# Transcription directs Holliday junction branch migration

**DOI:** 10.64898/2026.03.13.711646

**Authors:** Tom J. Powell, George G. B. Brown, Rachal M. Allison, Jon A. Harper, Matthew J. Neale, William H. Gittens

**Affiliations:** Genome Damage and Stability Centre, School of Life Sciences, University of Sussex, Brighton BN1 9RQ

## Abstract

During meiosis, genetic diversity arises from the resolution of branched DNA intermediates called Holliday junctions to create crossovers—sites of reciprocal exchange between parental chromosomes. Holliday junctions arise during the repair of Spo11-induced DNA breaks, yet the principles linking break formation, repair, and chromosome architecture remain unclear. Top3, a type IA topoisomerase, acts on Holliday junctions, but its spatiotemporal dynamics are unknown. Here, we map Top3 catalytic activity throughout meiotic prophase with strand specificity and nucleotide resolution. We identify a DNA sequence motif associated with catalysis and a pattern of activity around Spo11 hotspots that requires ongoing repair and Top3’s helicase partner, Sgs1. Strikingly, Top3 activity shifts over time, influenced by transcription, cohesin, and the crossover factors Msh5 and Mer3, redistributing toward sites of convergent transcription—known locations of meiotic cohesin association. Upon prophase exit, as crossovers are resolved, Top3 activity subsides. Remarkably, maps of genome-wide recombination reveal that crossover resolution preferentially occurs at these same regions of convergent transcription. Collectively, we propose that Top3 coordinates the transcription-coupled movement of Holliday junctions from Spo11 hotspots towards cohesin-associated axis sites, whereupon crossover resolution occurs to ensure accurate meiotic chromosome segregation.

## RESULTS

We previously demonstrated that rapid fixation enables the endogenous activity of Topoisomerase 2 (Top2) to be mapped with high resolution across the *S. cerevisiae* genome^1^. Because this mapping methodology (CC-seq; **Fig. S1a**) is specific for proteins covalently linked to DNA via a 5ʹ phospho-tyrosine bond (covalent complexes, CC, typified by topoisomerase-family enzymes) we reasoned that it may also reveal sites of Top3 activity, which, along with Spo11, form the same type of covalent protein–DNA bond^2,3^ (**Fig. 1a**). To explore this idea, we generated CC-seq maps from *S. cerevisiae* cells synchronised in meiotic prophase—an evolutionarily conserved process in which a high rate of genetic recombination occurs and where a central role for Top3 has been implicated^4-7^ (**Fig. 1a**).

**Figure 1.**
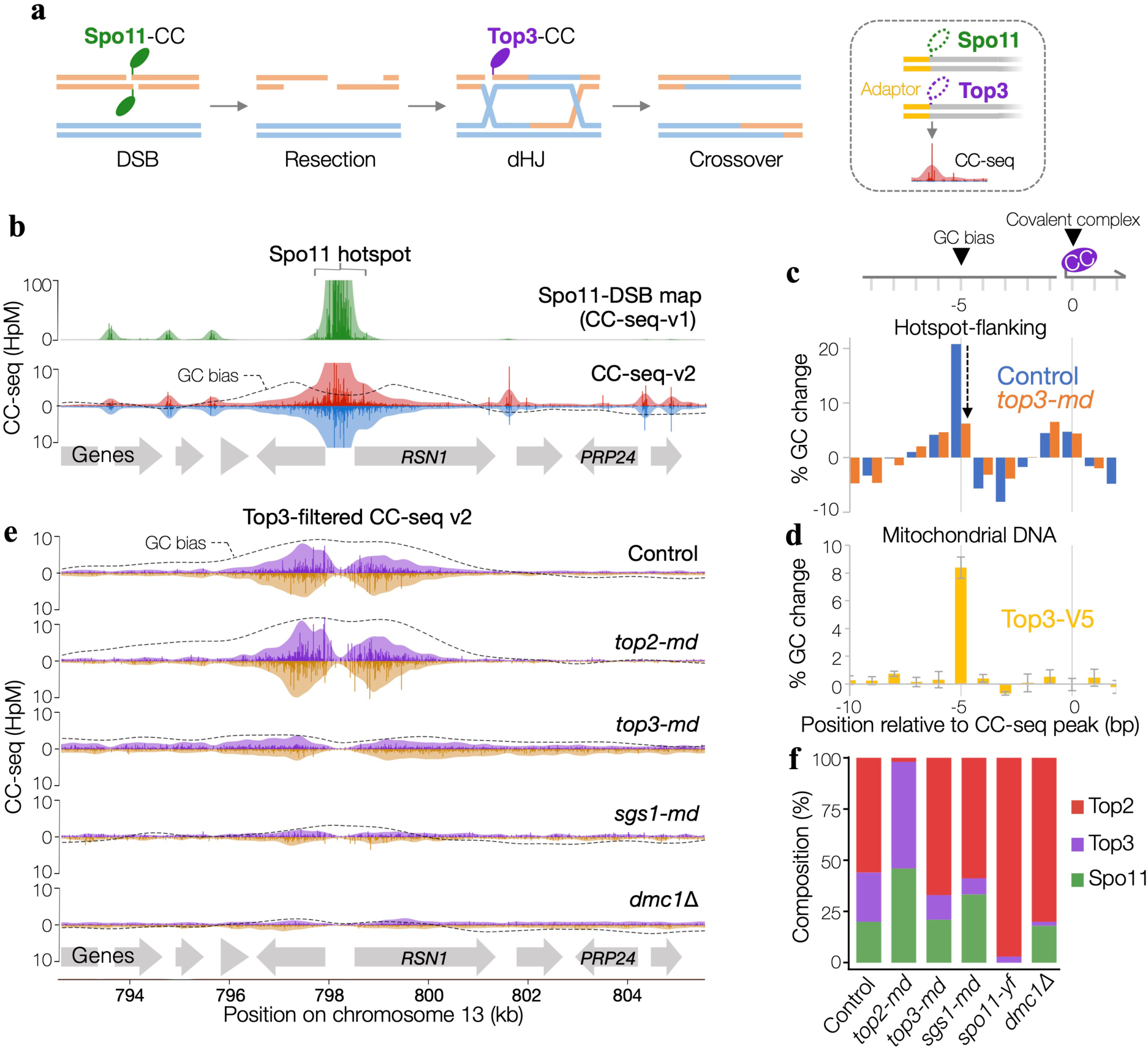
CC-seq maps Top3 activity. **a,** Cartoon of meiotic recombination pathway with Spo11 and Top3 covalent complexes indicated, which are enriched and mapped onto genomic coordinates (CC-seq). **b,** CC-seq maps over a 20 kb region around a Spo11-DSB hotspot. Green trace, Spo11-DSBs averaged over 2–4 hr in *ndt80*Δ *sae2*Δ cells. Red/blue tracks are unfiltered CC-seq-v2 maps of proteins covalently linked to DNA via 5′ phosphotyrosine bonds (Spo11, Top2, Top3) in control (*ndt80*Δ) cells. Red and blue traces denote top- and bottom-strand 5′ termini. **c,** Base composition around unfiltered CC-seq-v2 signal peaks in 500 bp windows flanking Spo11-DSB hotspots. Data show percentage change in GC content relative to the local average. **d,** Base composition around CC-seq signal peaks enriched by anti-V5 immunoprecipitation of Top3–V5 in mitochondria, shown as percentage change in GC content relative to a negative control immunoprecipitation of the same strain using protein G. **e,** Top3-filtered CC-seq-v2 maps (masked to remove Spo11 and Top2 signals) in control and mutant strains. Purple and gold traces denote top- and bottom-strand 5′ termini. Arrows indicate gene positions. **f,** Stacked bar plot showing library signal composition in control and mutant strains (all *ndt80*Δ). Top2 (red), Spo11 (green) and Top3 (purple) signal fractions represent the best-fitting combination of reference sequence-bias profiles identified by minimum root mean square deviation (RMSD) from the observed unfiltered signal (**Fig. S4a-b**). HpM, hits per million mapped reads. In all maps, lighter tones show data smoothed with a 500 bp sliding Hann window. Dashed lines indicate skews towards GC bases 5 bp upstream of cleavage (log_2_ axis range of ±4).

### Genome-wide maps of Topoisomerase 3 activity

In contrast to the Top2-enriched CC-seq maps generated from cells arrested in G1^1^, meiotic maps are anticipated to be a composite pattern of Top2, Spo11 DSBs, plus also putative Top3 signals associated with recombination. Comparing maps generated at 3 hours following meiotic induction—when recombination will be ongoing—to a sample enriched only for Spo11 DSBs (**Fig. 1b, upper**) revealed signals that spread either side of Spo11-DSB hotspot areas on both strands (**Fig. 1b, lower**). Additional narrow sporadic peaks were also present, which were absent in the ‘Spo11-DSB only’ control track (**Fig. 1b**) prepared from *sae2*Δ cells in which Spo11-linked DSBs accumulate and no recombination occurs.

To elucidate the origin of the various signals, the 3 h sample was compared to those in which either Top2 or Top3 were subjected to meiotic depletion. Whilst many of the narrow peaks were reduced by depleting Top2, the broad signal flanking Spo11 hotspots was not (**Fig. S1b-c**). By contrast, the hotspot-associated signal was reduced upon depletion of Top3, or Sgs1 (the helicase binding partner of Top3) (**Fig. S1c**), supporting the view that these flanking signals represent direct detection of Top3 covalent intermediates. Indeed, as expected from Top3’s known role in meiotic recombination, hotspot-flanking signals were dependent on both Spo11 catalytic activity^2,8^ (**Fig. S1c**), and DNA strand invasion mediated by the meiotic RecA/Rad51 orthologue Dmc1^9^ (**Fig. S1c**).

Notably, CC-seq signals were also enriched in the mitochondrial DNA (mtDNA) genome irrespective of Top2 depletion, but dependent on Top3 (**Fig. S2a–d**), suggesting they represent Top3 activity within mitochondria, where it has been previously implicated in other organisms^10-13^. Unlike hotspot-adjacent signals, mitochondrial Top3 activity was independent of Sgs1 (**Fig. S1a-b**)—in agreement with observations in human cells^11^.

Analysis of the base composition of the mtDNA signals revealed a strong skew towards G or C bases 5 bp upstream (hereafter -5GC) of the mapped 5′-linked covalent complex (**Fig. S2e-f, i-j**). Such a bias depended strongly on Top3 (**Fig. S2e-f, i-j**), and, notably, is supported by studies of yeast Top3 cleavage sites within a synthetic substrate in vitro^14^. Moreover, in the nuclear genome substantial Top3-dependent -5GC skews were observed in Spo11 hotspot-flanking regions (**Fig. 1b**, dashed line; **Fig. 1c** and **Fig. S2g**) that were independent of Top2 (**Fig. S1c**), but depleted upon loss of Top3, Sgs1, Spo11 or Dmc1 activity (**Fig. 1c**, **Fig. S1c** and **Fig. S2g**). Finally, CC-seq libraries generated from antibody-based enrichment of V5 epitope-tagged Top3 displayed a similar -5GC skew that was absent in the untagged Top3 control (**Fig. S2h**), and in a mock immunoprecipitation using nonspecific protein G (**Fig. 1d** and **Fig. S2h**). Taken together, these data further support the view that the enriched signals detected in vivo around Spo11 hotspots are maps of Top3 covalent intermediates, and thus indicative of Top3 catalytic activity—something never before characterised.

We next generated a filtered view of signals that we define as Top3 dependent. We defined high-confidence genome-wide maps of Spo11 and Top2 activity and used them as nucleotide-resolution masks to remove Spo11 and Top2 sites from the data (**Fig. S3a**). Whilst this approach is stringent—in that Top3 signals that exactly colocalize with Spo11 and Top2 will also be removed—importantly, 91% of the genomic space remained unmasked, and as such, mappable. In control cells, such masking abolished peaks previously ascribed to Top2 and Spo11, yet retained broad regions of enrichment on both strands around Spo11 hotspots at this mid-prophase (3 hour) timepoint that also displayed the -5GC bias (**Fig. 1e**). Equivalent maps generated upon depletion of Top2 were largely unchanged (**Fig. 1e**), whereas depletion of Top3 or Sgs1, or absence of Dmc1, reduced or abolished these features (**Fig. 1e**). To determine if -5GC was absolutely necessary for cleavage by Top3, masked data was subset based on presence of a -5GC or -5AT relative to each peak. Strikingly, hotspot-adjacent Top3 activity was almost entirely dependent on the presence of -5GC (**Fig. S3b**).

To estimate the global proportions of Spo11, Top2, and Top3 within the entire unfiltered 3-hour datasets, we exploited the characteristic DNA sequence skews associated with each class of covalent complex (**Fig. S4a**) to decompose the mixture into its component fractions (**Fig. 1f** and **Fig. S4a-b**). In control cells, about half the library was composed of Top2, with the remaining proportions of Top3 and Spo11 roughly equal (**Fig. 1f**). As expected, Top2 depletion largely removed all Top2 signal without any significant changes in the relative proportions of Spo11 and Top3 (**Fig. 1f**). By contrast, depletion of Top3 itself (known to be incomplete^7^), and to a greater extent deletion of Sgs1, substantially reduced the Top3 activity detected (**Fig. 1f**). Finally, consistent with the filtered maps (**Fig. 1e**), Top3 signals were largely undetectable in the absence of either Spo11 or Dmc1 activity (**Fig. 1f**).

Collectively, these observations indicate that our maps and filtering methodology enable the detection and characterisation of spatial patterns of Top3 activity associated with meiotic recombination.

### Top3 activity migrates codirectionally with local transcription towards chromosome axis sites

We next sampled timepoints through meiotic prophase in synchronised cells that were run into a permanent block in late prophase mediated via deletion of the *NDT80* transcription factor^15^ (**Fig. 2a**), where recombination intermediates are known to accumulate^16^. Remarkably, Top3 signals displayed clear spatiotemporal changes—being found in hotspot-flanking regions at 3 hours, but spreading more widely, and often accumulating in strong peaks displaced 1–3 kb from hotspots by 4–6 hours (**Fig. 2a**). Visual inspection revealed a clear association between peaks of Top3 activity at late timepoints and sites of convergent transcription (**Fig. 2a**). Such chromosomal sites are preferentially bound by meiotic cohesin (mapped by ChIP-seq of the meiotic kleisin subunit Rec8^17^; **Fig. 2a**), which establishes chromatin loops and organises recruitment of axial proteins^18,19^.

**Figure 2.**
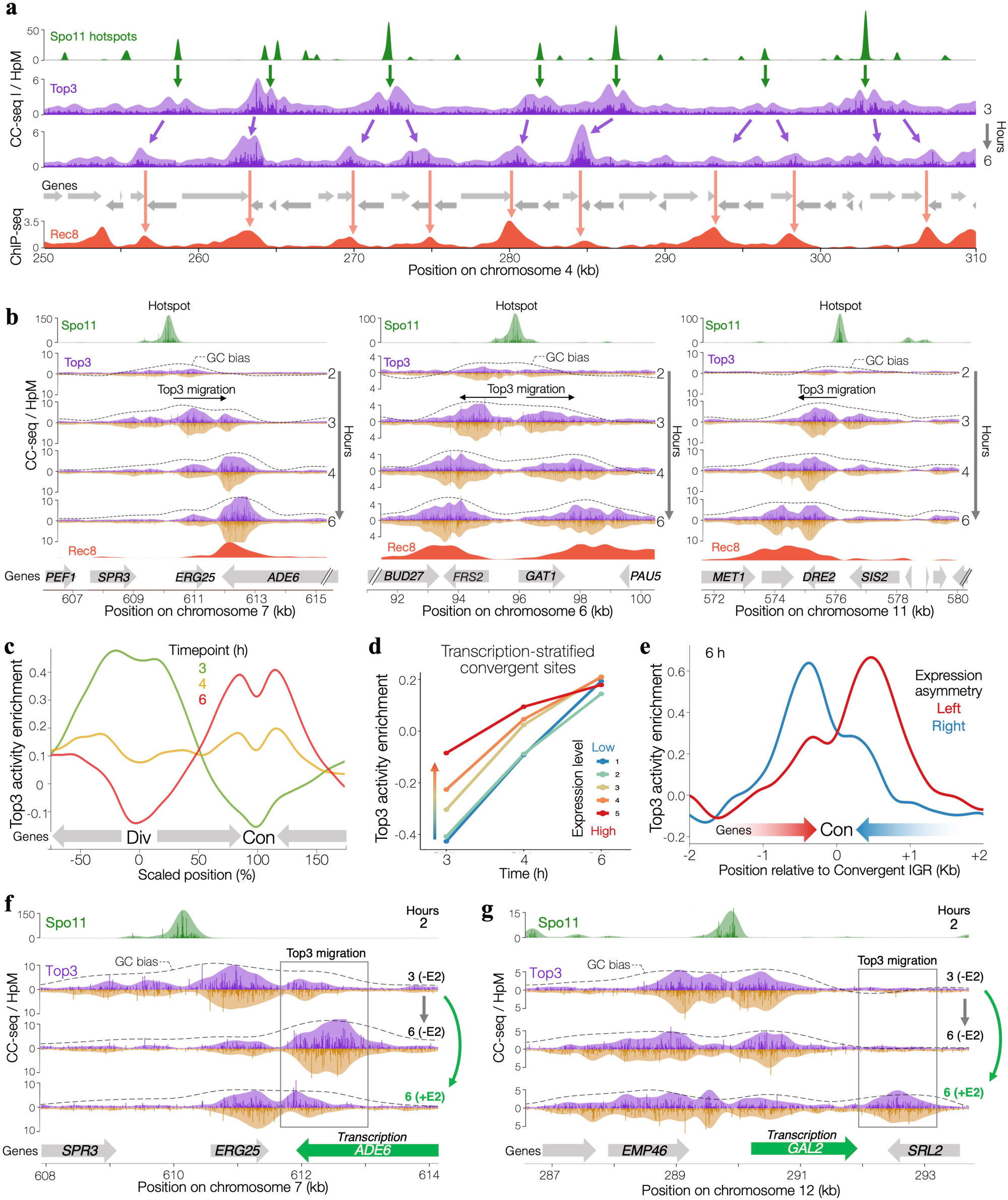
Top3 activity tracks transcription towards chromosome axis sites. **a,** Composite maps of Spo11 hotspots (CC-seq v1, 2–4 h, *ndt80Δ sae2Δ*, brown), Top3 activity (Top3-filtered CC-seq v2, 3 h and 6 h, *ndt80Δ*, purple), and Rec8 occupancy (ChIP-seq, 3 h, orange). Grey arrows mark genes; brown arrows indicate early Top3 overlap with Spo11 hotspots, purple arrows Top3 migration, and orange arrows colocalization with Rec8 peaks. **b,** Expanded views of three 9 kb regions showing strand-specific Spo11 (brown/green) and Top3 (purple/gold) activity with Rec8 occupancy (orange). Lighter tones show 500 bp Hann smoothing; dashed lines mark skews towards GC bases 5 bp upstream of cleavage (log_2_ axis range of ±4). **c,** Average Top3 activity enrichment across scaled intervals between 1,334 neighbouring divergent and convergent intergenic regions (IGRs). **d,** Median Top3 enrichment within 1 kb windows centred on convergent IGRs, stratified by summed convergent transcription, at 3, 4, and 6 hours after meiotic induction in *ndt80*Δ cells. **e,** Average Top3 activity enrichment around convergent IGRs, separated by transcriptional asymmetry (top 10% left-biased, red; right-biased, blue). In **c-e** enrichment is the ratio of Top3-filtered CC-seq v2 over local mappability (100 bp Hann smoothing). **f-g,** Composite maps at estradiol-inducible loci (*P_GAL1_ADE6* and *GAL2*) in *ndt80Δ GAL4::ER P_GAL1_ADE6 P_GAL1_KIN1* cells. Top3 activity (purple/gold, 3 hours meiosis induction ±3 hr estradiol induction), and reference track of Spo11 activity (green) are shown with 500 bp Hann smoothing. Dashed lines mark -5GC skews; grey arrows mark genes; green arrows indicate induced transcription. HpM, hits per million mapped reads.

At fine scale, Top3 activity that was initially associated with Spo11 hotspots migrated in a pattern correlated with the local direction of transcription (**Fig. 2b** and **Fig. S5a-c**). Such migration typically spanned 1–4 kb, before accumulating at, or close to, sites of convergent transcription—a trend we confirmed globally by aggregating Top3 dynamics across scaled intervals between divergent and convergent intergenic regions (IGRs; **Fig. 2c**).

Quantitative links between the rates of local transcription and Top3 migration were tested by stratifying loci based on the combined expression level of converging gene pairs (**Fig. 2d**). At the earliest timepoint (3 hours), transcription and Top3 were positively correlated, with more highly transcribed convergent loci containing more Top3 activity. At late timepoints, this trend weakened, such that by 6 hours all convergent loci had similarly high levels of Top3 activity (**Fig. 2d**). This relationship suggests that the rate of Top3 migration is directly influenced by the rate of local transcription, with higher rates of transcription causing faster—and therefore earlier—migration of Top3.

Although convergent sites generally became enriched for Top3 activity over time, localised Top3 peaks were often displaced towards the 3′ end of one gene rather than centred directly over the convergent IGR (**Fig. 2b**). Similar patterns of Red1 occupancy, an important axial factor involved in DSB and crossover regulation, have been reported to be driven by relative differences in the expression rate of converging gene pairs^20^. Indeed, stratifying convergent loci by the relative strengths of converging gene pairs revealed Top3 to be patterned similarly, with enrichment in the 3′ end of the lesser-transcribed gene of the pair (**Fig. 2e**).

The causal influence of transcription was tested by placing two genes that faced towards sites of focal Top3 enrichment under the control of the *GAL1* promoter (**Fig. 2f** and **S6a-f**). Without transcriptional induction—where expression levels are similar to that of the native genes (**Fig. S6a-b**)—Top3 spread from the hotspot and accumulated at the convergent sites similarly to control cells (**Fig. 2f** and **Fig. S6d-e**). By contrast, inducing transcription at 3 hours reduced the spreading towards, and accumulation of Top3 within, the extreme 3′ terminus of the induced *ADE6* gene (**Fig. 2f** and **Fig. S6d**), but little effect at *KIN1* in which Top3 signals continued to accumulate beyond the end of the annotated *KIN1* reading frame (**Fig. S6e;** Discussion).

We next asked whether migration could be promoted by inducing tandem (concurrent) transcription, by examining the native *GAL2* locus, whose promoter harbours a Spo11 hotspot (**Fig. 2g** and **Fig. S6f**). In the absence of induction, Top3 signals were impeded by *GAL2*, and remained proximal to the Spo11 hotspot (**Fig. 2g** and **Fig. S6f**). Strikingly transcriptional activation resulted in a substantial fraction of the Top3 signal migrating ∼2 kb through *GAL2* and accumulating within the 3′ region of the convergent *SRL2* gene (**Fig. 2g** and **Fig. S6f**). Collectively, these observations demonstrate that transcription has the potential to rapidly pattern the migration of recombination-dependent Top3 activity.

### Top3 accumulation at axis sites depends on cohesin

Cohesin organises chromosomes into longitudinally compacted linear arrays of loops^18,19,21,22^, potentially via its loop-extruding motor activity^23,24^—something that may work alongside and/or antagonistically with transcription^25^. Because sites of convergent transcription are also locations in which the occupancy of the cohesin subunit, Rec8, is greater^26^, there was also a positive association between Top3 migration and Rec8 abundance as detected by ChIP-seq (**Fig. 2a-b**).

To explore the influence of cohesin on Top3 spatiotemporal patterning, we deleted *REC8* and compared CC-seq maps to controls (**Fig. 3**). *REC8* deletion leads to a redistribution of Spo11 activity into areas of the genome referred to as ‘islands’ (**Fig. S7a-b**^27^). As expected, given its dependency on Spo11-DSB formation, we similarly saw a bias of Top3 signals to island regions (**Fig. S7a,c-d**). In island regions, *REC8* deletion reduces the accumulation of Top3 activity at convergent sites over time (**Fig. 3a** and **Fig. S5a-c**), a consistent trend observed when aggregating signals across scaled divergent–convergent intervals (**Fig. 3b)**. Aggregating Top3 signals across Rec8-bound sites revealed that greater Rec8 occupancy positively correlates with greater Top3 activity at 6 h in control cells (**Fig. 3c**). By contrast, in Rec8-deficient cells, these same sites showed reduced Top3 activity and a negative correlation (**Fig. 3d**). Globally, *REC8* deletion also displayed a delay in the onset of Top3 activity by at least one hour (**Fig. 3e** and **Fig. S7d**) consistent with Rec8, and therefore cohesin, being important for efficient meiotic recombination^22^. We conclude that at least part of what patterns Top3 activity is the loop-forming activity of cohesin.

**Figure 3.**
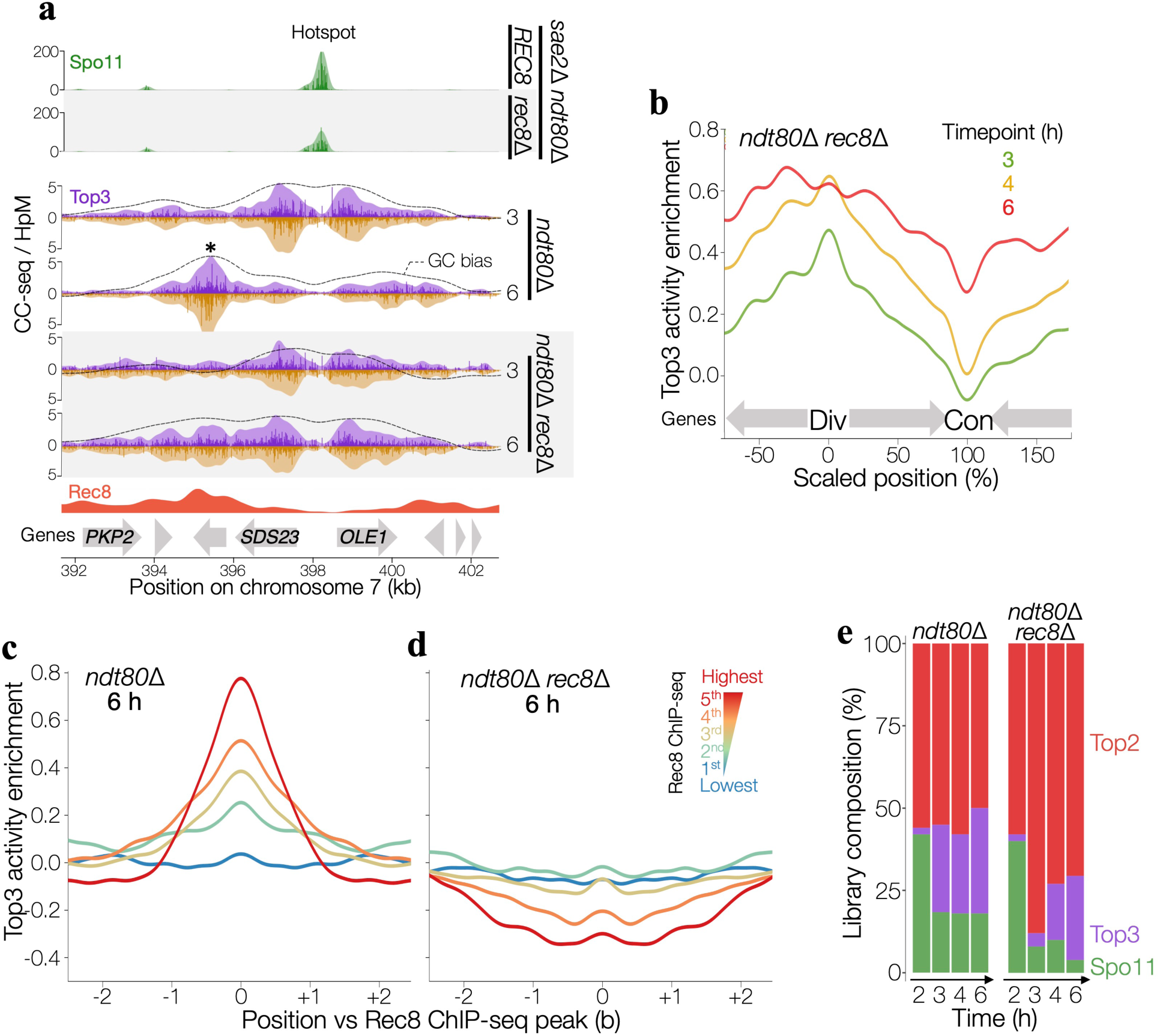
Top3 accumulation at axis sites depends on cohesin. **a**, Composite maps of strand-specific Spo11 activity (CC-seq v1, 2–4 h, *ndt80Δ sae2Δ* and *ndt80Δ sae2Δ rec8*Δ, green), Top3 activity (Top3-filtered CC-seq v2, 3 h and 6 h, *ndt80Δ* and *ndt80Δ rec8*Δ, purple/gold), and Rec8 occupancy (ChIP-seq, 3 h, orange), at a ∼10 kb region. Lighter tones show 500 bp Hann smoothing; dashed lines mark skews towards GC bases 5 bp upstream of cleavage (log_2_ axis range of ±4). Grey arrows mark genes; asterisks highlight accumulation points diminished in *ndt80Δ rec8*Δ. **b**, Average Top3 activity enrichment across scaled intervals between 191 neighbouring divergent and convergent intergenic regions (IGRs) within island regions. HpM, hits per million mapped reads. **c-d,** Average Top3 activity enrichment in *ndt80Δ* (**c**) and *ndt80Δ rec8*Δ strains (**d**), around 2886 Rec8 binding sites stratified by the Rec8 ChIP-seq occupancy in wild-type Rec8 cells^20^. Enrichment is shown as a fold of Top3-filtered CC-seq v2 over local mappability (100 bp Hann smoothing). **e**, Stacked bar showing best-fit estimates for the relative fraction of Top2 (red), Spo11 (green) and Top3 (purple) signal at each timepoint (see Methods).

### Top3 migration towards axis sites depends on Class I CO factors Mer3 and Msh5

The migration of Top3 signals, and dependency on Spo11 and Dmc1, suggests Top3 activity is associated with the migration of a homologous recombination intermediate. Top3, as part of the STR (Sgs1-Top3-Rmi1) complex, has been implicated in both pro-crossover (CO) and pro-noncrossover (NCO) roles^6,7^. We tested for a role in CO formation via deletion of two of the known factors involved: Msh5 (part of the Msh4–Msh5 heterodimer, proposed to be involved in protection of early recombination intermediates^28,29^), and Mer3 (a DNA helicase proposed to stabilize and extend strand invasions in the 3′-5′ direction, thereby promoting the formation of double Holliday Junctions^30,31^). Remarkably, deletion of either factor almost completely abolished Top3 migration, with signals remaining associated with hotspot-flanking regions even after 6 hours (**Fig. 4a-b** and **S5a-c**)—a pattern recapitulated genome-wide when aggregated across scaled divergent–convergent IGR intervals (**Fig. 4c**). Such findings directly connect migration of Top3 activity to the pro-CO pathway, and suggest that CO-designated recombination intermediates (upon which Top3 is acting) are changing position during meiotic prophase.

**Figure 4.**
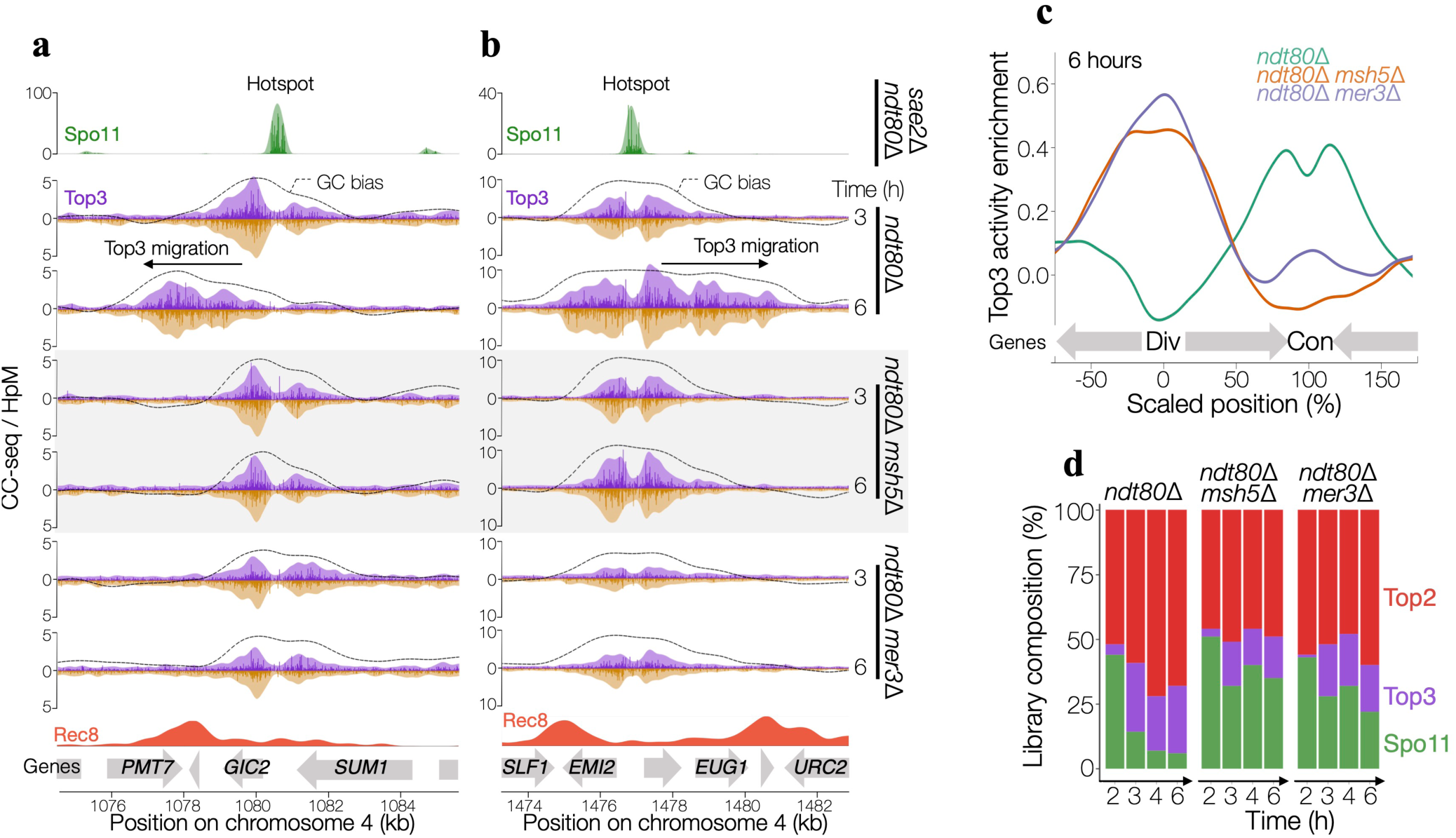
Top3 migration towards axis sites depends on Class I CO factors Mer3 and Msh5. **a-b**, Composite maps of strand specific Spo11 activity (CC-seq v1, 2–4 h, *ndt80Δ sae2Δ*, brown/green), Top3 activity (Top3-filtered CC-seq v2; 3 h and 6 h; *ndt80Δ, ndt80Δ msh5*Δ, *ndt80Δ mer3*Δ, and *ndt80Δ rec8*Δ; purple/gold), and Rec8 occupancy (ChIP-seq, 3 h, orange), at two ∼10 kb regions. Lighter tones show 500 bp Hann smoothing; dashed lines mark skews towards GC bases 5 bp upstream of cleavage (log_2_ axis range of ±4). Grey arrows mark genes. **c**, Average Top3 activity enrichment across scaled intervals between 1,334 neighbouring divergent (Div) and convergent (Con) intergenic regions (IGRs), at 6 hours following meiotic induction in control and mutant strains. Enrichment is shown as a fold of Top3-filtered CC-seq v2 over local mappability (100 bp Hann smoothing). HpM, hits per million mapped reads. **d**, Stacked bar showing best-fit estimates for the relative fraction of Top2 (red), Spo11 (green) and Top3 (purple) signal at each timepoint (see Methods), after removing end-adjacent regions (EARs).

The persistence of hotspot-adjacent Top3 activity in *mer3*Δ or *msh5*Δ cells could reflect either long-lived recombination intermediates or the formation of new substrates due to sustained Spo11 activity. To distinguish between these possibilities, we again decomposed the unfiltered libraries into individual signal components. In control cells, the minor amount of Spo11 signal that persisted at late time points (**Fig. 3e** and **Fig. S4c**) was enriched only near chromosome ends (end-adjacent regions, EARs^32^) and diminished when these regions were excluded (**Fig. 4d** and **Fig. S4d**). By contrast, in *mer3*Δ or *msh5*Δ cells, Spo11 signals persisted at substantially higher levels across the genome (**Fig. 4d** and **Fig. S4d**), suggesting that the hotspot-adjacent Top3 activity in these mutants arises from ongoing cycles of DSB formation and repair.

### CO resolution sites become associated with chromosome axis sites

To understand exactly how patterns of Top3 migration may correspond to patterns of downstream COs, we generated CC-seq libraries at multiple timepoints from wild-type cells, where CO-designated recombination intermediates are known to be transient^16,33,34^. As anticipated, and unlike in *ndt80*Δ-arrested cells, Top3 signals no longer accumulated at the 6 hour timepoint (**Fig. 5a** and **S4a-c**)—consistent with ongoing CO resolution removing those molecular intermediates upon which Top3 had been acting. Importantly, however, loci still displayed clear evidence of Top3 migration away from hotspots in the 3 and 4 hour timepoints (i.e. prior to resolution), often with partial, transient, enrichment at axial sites, similar to the *ndt80*Δ-arrested strain (**Fig. 5a** and **Fig. S5a-c**). Furthermore, at 3 h—when Top3 maps most closely resembled *ndt80*Δ maps—Top3 activity at convergent sites was positively associated with the level of convergent transcription (**Fig. 5b**), and its migration through genes correlated with the transcriptional level of those genes (**Fig. 5c**).

**Figure 5.**
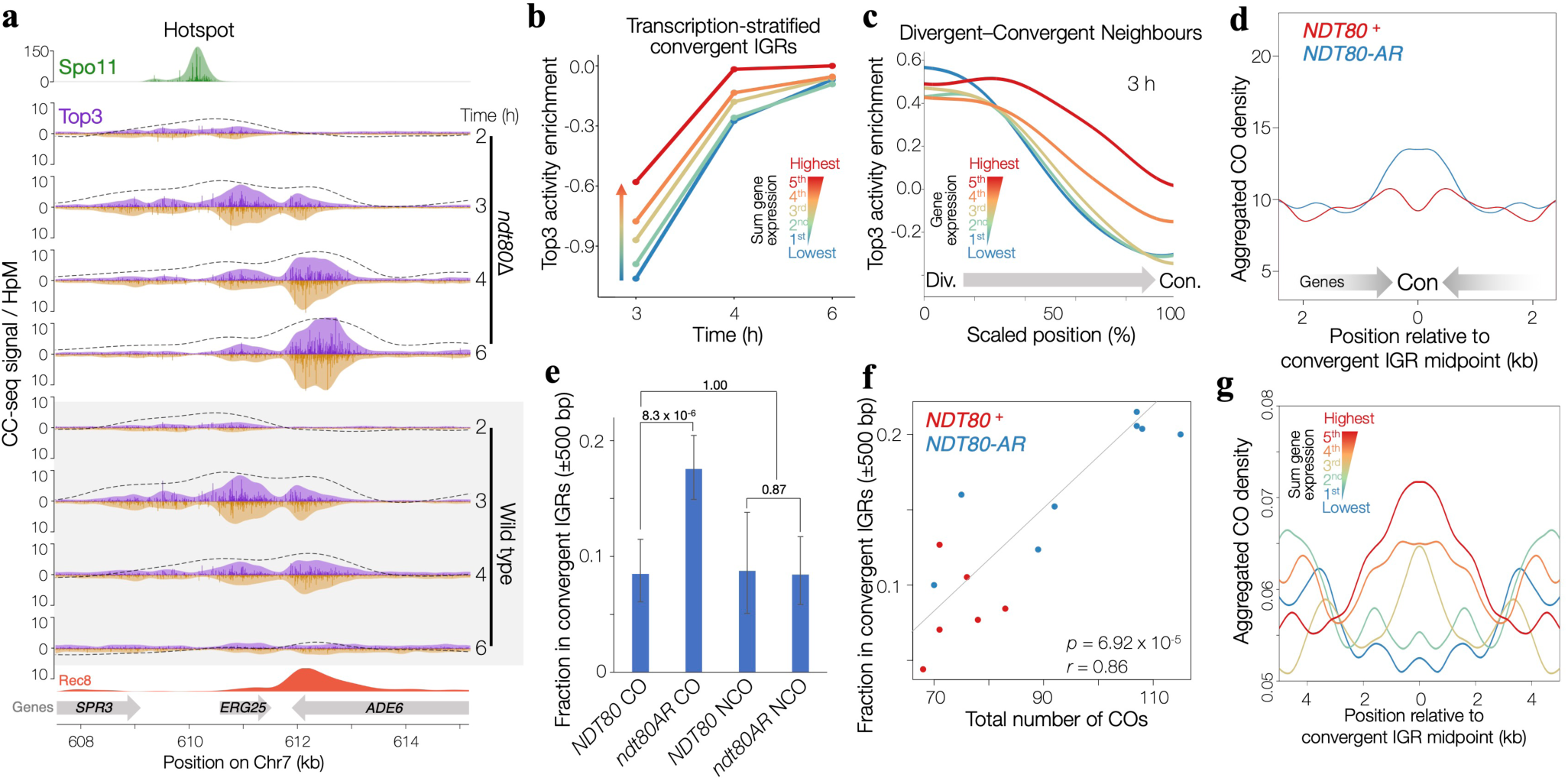
CO resolution sites become associated with axis sites over time. **a,** Composite maps of strand specific Spo11 activity (CC-seq v1, 2–4 h, *ndt80Δ sae2Δ*, brown/green), Top3 activity (Top3-filtered CC-seq v2; 3, 4, and 6 h; *ndt80Δ* and wild type, purple/gold), and Rec8 occupancy (ChIP-seq, 3 h, orange). Lighter tones show 500 bp Hann smoothing; dashed lines mark skews towards GC bases 5 bp upstream of cleavage (log_2_ axis range of ±4). Grey arrows mark genes. **b**, Median Top3 enrichment within 1 kb windows centred on convergent intergenic regions (IGRs), stratified by summed convergent transcription, at 3, 4, and 6 hours after meiotic induction in wild-type cells. **c**, Average Top3 activity enrichment across scaled intervals between 1076 neighbouring divergent (Div) and convergent (Con) intergenic regions (IGRs), at 6 hours following meiotic induction in wild-type cells, stratified by the expression level of the intermediary gene. Enrichment as fold of Top3-filtered CC-seq v2 over local mappability (100 bp Hann smoothing). **d**, Average crossover density around convergent intergenic regions (IGRs) pooled over 6 wild-type vs 8 *NDT80*-arrest-release (*NDT80-AR*) meioses. **e**, Proportion of total crossovers (COs) and noncrossovers (NCOs) found in convergent intergenic regions ±500 bp, in wild-type and *NDT80-AR* meioses. Bars 1–4 represent proportions ±95% binomial CIs for 447, 763, 183 and 379 total events, respectively, pooled over multiple meioses; *p*-values determined by Fisher’s exact test (two-sided). **f**, Total number of COs vs the proportion found in convergent intergenic regions ±500 bp, across 6 wild-type and 8 *NDT80-AR* meioses. The Pearson correlation coefficient (*r*) and associated *p*-value for the combined distribution are inset. **g**, Average CO density around convergent intergenic regions (IGRs) stratified by sum convergent transcription in 52 wild-type meioses^37,38^.

Next, we interrogated maps of CO formation previously collected from individual meiotic cells with and without transient extension of meiotic prophase^35^ mediated via temporal regulation of the *NDT80* gene^36^ (Ndt80 arrest-release, *ndt80AR*). Remarkably, aggregation of CO positions revealed an enrichment at sites of convergent transcription—axis sites—within cells that had been subjected to prophase extension (**Fig. 5d-e**; *P*-value = 8.3 x 10^-6^)—something that was not observed for NCOs (**Fig. 5e**; *P* = 0.87). Moreover, across individual meioses, there was a strong positive correlation (R = 0.86, *P* = 6.9x10^-5^) between total CO number and the proportion found within axis sites (**Fig. 5f**). Because each meiotic cell may spend a different length of time in prophase, and because a longer time in prophase is expected to be correlated with increased CO number^35^, these data directly link prophase length to alterations in CO positioning.

If Top3 migration is indeed linked to CO positioning, we hypothesised that CO distribution would similarly reflect transcriptional activity. To test this, we analysed a large dataset that collectively mapped 4783 COs in 52 wild-type meioses^37,38^. Remarkably, CO positioning showed a positive association with convergent transcription: COs were depleted at weakly expressed convergent sites and enriched at highly expressed convergent sites (**Fig. 5g**).

Collectively, these observations indicate that local transcription patterns movement of CO-designated recombination intermediates, upon which Top3 acts, towards sites of convergent transcription—known to be preferred sites of chromosome axis association—suggesting that CO resolution may occur favourably when CO-designated precursors are spatially associated with the chromosome axis.

## DISCUSSION

Chiasmata provide the physical linkages that ensure biorientation and accurate segregation of homologous chromosomes during meiosis I. The establishment of these linkages requires tight coordination between the nanoscale molecular events of DNA recombination and microscale structural rearrangements of meiotic chromosomes, including synaptonemal complex dynamics and remodelling of homolog axes. Although factors such as the ZMM proteins have been identified at the interface of these processes^39,40^, the spatiotemporal dynamics that couple recombination with chromosome-axis morphogenesis remain poorly understood.

In *S. cerevisiae*, Spo11-induced DSBs predominantly occur in promoter regions, within chromatin loops (**Fig. 6a**), whereas much of the protein machinery required for their formation and processing is concentrated at the axis—organised around cohesin at sites of convergent transcription. This spatial disconnect has led to the proposal that protein–protein interactions tether DSB sites to the axis. Yet how such potential loop–axis tethering influences downstream recombination steps remains unclear. For example, are tethered DSBs released to facilitate homology search, and if so, at what stage? Is the invaded DNA loop also tethered to its axis, and if so, how is this coordinated?

**Figure 6.**
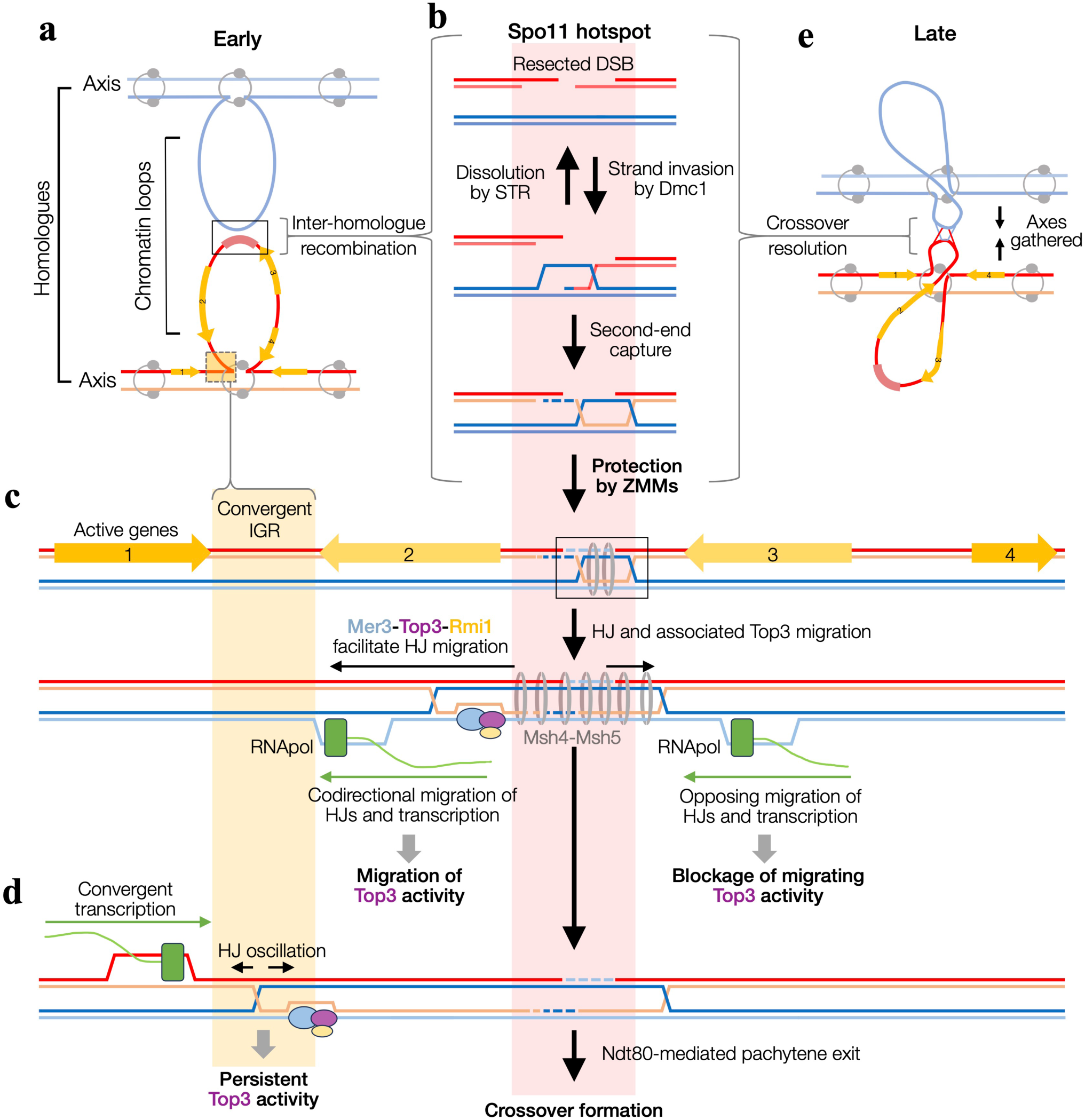
Transcription directs Holliday junction branch migration. **a,** Simplified meiotic loop–axis structure of homologous chromosomes (red and blue lines). Cohesin (grey rings) links sister chromatids (pale and dark blue/red), and organizes intrachromosomal loop bases (only one shown for clarity) at sites of convergent transcription. **b,** Spo11-DSBs form at hotspot regions within chromosome loops. Following resection, strand invasion generates early recombination intermediates that can be dismantled by Sgs1–Top3–Rmi1 (STR) or proceed through second-end capture and ZMM-mediated protection toward crossover-designated double Holliday junctions (dHJs). **c**, We propose that Mer3–Top3–Rmi1 facilitates Holliday junction (HJ) branch migration, resulting in relocation of Top3 activity away from the initiating hotspot along chromatin. Productive migration is favoured when codirectional with transcription and impeded when opposing the direction of transcription. **d**, Convergent transcriptional sites, which are also preferential loop bases bound by cohesin, act as barriers that stall migrating junctions, causing persistent Top3 activity that we propose is associated with local HJ oscillation. **e,** The migration of HJs towards loop bases will tend to (1) spatially gather homologous chromosomes and (2) concentrate sites of HJ resolution and thus CO formation with the chromosome axis.

By directly mapping Top3 catalytic activity around Spo11 hotspots, we corroborate—and extend—genetic and biochemical evidence for a role of Top3 in meiotic recombination. The spatiotemporal dynamics and genetic dependencies of these maps indicate that Top3 acts on early recombination intermediates at hotspots, both near the DNA break and upon a subset of intermediates that migrate away (**Fig. 6b-d**)—a hallmark of CO formation^41^. Notably, helicase-driven migration of double Holliday junctions (dHJs) requires Top3 activity^42,43^. Although Top3–Rmi1 is classically partnered with Sgs1 in the STR complex to disassemble joint molecules, recent work suggests that Mer3 can compete with Sgs1 for Top3–Rmi1, defining an alternative Mer3–Top3–Rmi1 (MTR) assembly^44^. Importantly, Mer3—cooperatively with the MutSγ heterodimer (Msh4-Msh5)—is known to stabilise joint molecules and promote their eventual repair as COs^28,30^.

Consistent with this, the Mer3 and Msh5 dependence of the migrating Top3 signal fraction supports a model in which MTR-MutSγ ensemble promotes the movement of single-end invasions (SEIs) and/or Holliday junctions (HJs) away from hotspots, thereby stabilizing them for downstream processing as COs (**Fig. 6c-d**). This model could explain why Mer3 is associated with both Spo11-DSB hotspot regions and axial sites *in vivo*^45^, which has been previously ascribed to the transient interactions between DSB and axis sites^17^.

Our observations reinforce the concept that branch migration stabilizes and helps designate recombination intermediates for CO resolution^41^. We further propose that such migration also moves essential recombination intermediates into closer proximity with the meiotic axis—and causes spatial gathering of homologous chromosomes (**Fig. 6e**). Thus the designation of COs becomes spatially and mechanistically linked to the chromosome axis, where downstream remodelling events (synaptonemal complex formation, axis reorganisation) associated with the essential process of chiasmata formation can be triggered^39,40^.

In the absence of Mer3, Top3 activity persists proximal to hotspots, consistent with rapid STR-dominated disassembly. Such persistence is readily explained by iterative cycles of D-loop dismantling and reinvasion^41,46^, many of which culminate in synthesis-dependent strand annealing to yield noncrossovers^16,39,40^. In the absence of CO formation, Spo11 DSBs may continue to form^39,40,47^, fuelling additional turnover by STR. Supporting this, at late time points Spo11-DSB signals persist in *mer3*Δ and *msh5*Δ cells (**Fig. S4c-d**). By contrast, in ZMM-proficient cells, it is the migrated Top3 activity that persists, suggesting that displaced recombination intermediates remain dynamic and can undergo limited bidirectional oscillation, which may be functionally important for maintaining HJ architecture in the context of transcriptional forces and loop–axis mechanics. Consistent with this, a very recent study shows that Top3 is required to maintain dHJs and prevent their conversion into an aberrant species that displays extremely low electrophoretic mobility, indicative of a highly branched or entangled topology^48^. By revealing that late-prophase Top3 activity occurs at the recombination–transcription interface, our data support the idea that, without functional Top3, it is these encounters that can become actively damaging to dHJ architecture.

Intriguingly, we observe that Top3 migration is highly correlated with orientation and transcription rate of nearby genes. This could be the case if transcription-driven supercoiling, and/or locomotive RNA polymerase complexes directly pattern the migration of recombination intermediates. Supporting this idea, not only are constrained Holliday junctions a barrier to transcriptional elongation^49^, but when unconstrained, HJs are able to migrate in response to torsion^50^. Our results clearly demonstrate that transcription is ongoing *in cis* with meiotic DSB repair, and poses further questions. For example, how do the forces generated by transcription interplay with those generated by Sgs1 and Mer3 helicases? Additionally, given cohesin is also patterned by ongoing transcription—possibly via a loop extrusion model^23-25^—is branch migration coupled to this process? Whatever the precise mechanisms at play, it is notable that Top3 activity was not influenced by transcriptional changes at *KIN1* (unlike at *ADE6*), suggesting that effects are extremely local and thus unlikely to be mediated by long-range forces.

Type IA topoisomerases, exemplified by Top3, are universally conserved. Yet, owing to the absence of molecular strategies to capture their transient cleavage intermediates, genome-wide maps of their catalytic activity have lagged those of type IB (Top1) and type IIA (Top2) enzymes. By employing our updated CC-seq methodology—recently applied to map transient Top2 cleavage complexes^1^—we have determined the first genome-wide maps of Top3 activity, paving the way for dissecting Top3 function in diverse biological contexts, including chromosome segregation, DNA replication and transcription, and DNA repair outside of meiosis. Remarkably, Top3 cleavage sites are invariably positioned five nucleotides 3′ of guanine or cytosine residues, consistent with assays of purified yeast Top3 on synthetic substrates in vitro^14^. By contrast, comparable in vitro analyses of bacterial TopoIII and human Top3α/Top3β identified a strong preference for cytosine at this position, with little or no enrichment for guanine reported^51-53^. This discrepancy may reflect genuine species-specific differences in sequence preference, or technical limitations of the in vitro datasets, including the small number of sites and substrate contexts sampled. Notably, our preliminary analysis of candidate Top3α signals in human mitochondria suggests that human Top3α also accommodates both cytosine and guanine at the −5 position in vivo. Recent structural analysis of human Top3β in complex with a model ssDNA substrate indicate that sequence specificity is likely conferred by multiple hydrogen bonding and salt bridge interactions with the base at the -5 position^54^.

In summary, our elucidation of Top3 activity on recombination intermediates supports a unifying model in which DNA recombination events that initiate within chromatin loops may be conveyed towards distal chromosome axis sites via branch migration of HJs in a transcriptionally co-orientated manner. By driving the spatial convergence of recombination and axis sites (**Fig. 6e**), this mechanism could coordinate the molecular development of crossovers with the architectural maturation of chromosomes—accommodating a role for transcriptional and/or topological forces in the physical choreography of meiotic recombination.

## METHODS

### Strains

For meiotic experiments, strains are diploid isogenic derivatives of the SK1 strain background^55^. For mitotic experiments, strains are haploid isogenic derivatives of the W303 background. Strains were made by standard mating and/or transformation. A full strain list is provided in **Table S1**. During this study, we found that three strains (GB42, LH221, TP57) were heterozygous for a natural *PPH3* variant. We therefore regenerated these strains with homozygous wild-type *PPH3* (TP131, TP128, WG165); no differences were detected in the phenotypes tested, so data from both versions were combined (averaged).

### Synchronous timecourses

Synchronised meiotic cultures were prepared using the *P_CUP1_-IME1* system^56^. Briefly, diploid colonies were pre-grown in 4 mL YPD (1% yeast extract, 2% peptone, 2% glucose supplemented with 0.5 mM adenine and 0.4 mM uracil) overnight, diluted to OD600=0.3 in 150 mL BYTA (1% yeast extract, 2% tryptone, 1% potassium acetate, 50 mM potassium phthalate, 0.001% antifoam 204) and grown for 16 hours with rapid shaking until OD600≈17. 250 mL of fresh SPM (1% potassium acetate; 0.001% antifoam 204; supplemented with 5 µg/ml of adenine, arginine, histidine, tryptophan and uracil; and with 15 µg/ml leucine) was inoculated to OD600=2.5, and incubated at 30 °C with vigorous shaking. After 2 hours, CuSO4 was added to a final concentration of 50 µM, and referred to as the 0 minute timepoint. Samples were collected at intervals to create CC-seq-V2 libraries.

### CC-seq library preparation

CC-seq library preparation was performed as described previously^1,57^. To generate mixed maps of stable Spo11-DSBs and transient Top2 and Top3 covalent complexes (CCs), cells were fixed by direct addition of culture to -20°C ethanol (final concentration 70%), rapid mixing, and storage at -20°C until future use (CC-seq v2). Alternatively, to generate maps highly-enriched for stable Spo11-DSBs^57^, cell culture was chilled in ice water for 5 minutes, harvested by centrifugation without prior ethanol fixation, and pellets stored at -20°C until future use (CC-seq v1)^58^. In both cases, cells were spheroplasted with Zymolyase 100T (AMS Bio), fixed by mixing with 2 volumes of -20°C ethanol, pelleted and lysed with STE buffer (2% SDS, 0.5 M Tris pH 8.1, 10 mM EDTA, 0.05% bromophenol blue). Lysates were extracted with Phenol/Chloroform/IAA extraction (25:24:1 ratio) and the aqueous phase was precipitated with ethanol, washed, dried and then dissolved overnight in TE (10mM Tris Base.HCl pH 8.1, 1 mM EDTA) at 4°C. DNA was fragmented to an average size of 500 bp using a Covaris M220 (duty cycle 10%, peak power 75 W, 200 cycles/burst, 7°C, 24 min)—validated by agarose electrophoresis of a small sample—before adjusting buffers to column binding conditions (0.3 M NaCl, 0.2% Triton X-100, 0.1% N-lauroylsarcosine). CCs were enriched on QIAquick silica columns via centrifugation, washed TEN buffer (10 mM Tris-HCl pH 7.5, 1 mM EDTA, 0.3 M NaCl), and eluted with TES (10 mM Tris-HCl pH 7.5, 1 mM EDTA, 0.5% SDS). Eluates were Proteinase K treated and ethanol precipitated, then resuspended in TE and quantified by Qubit.

End repair and first adapter ligation (sonicated end) were performed with NEBNext Ultra II using a custom P7 adapter^58^. Unligated ends were dephosphorylated with alkaline phosphatase (rSAP, NEB), followed by cleanup with AMPure XP beads. Samples were treated with recombinant TDP2 to resolve 5′-phosphotyrosyl linkages^1,57,59^, cleaned up using AMPure beads, and subjected to a second ligation with a custom dephosphorylated-P5 adapter without prior end repair. Following cleanup and quantification, libraries were PCR-amplified with NEBNext Ultra II (dual index primers), cleaned up using AMpure beads, and assessed by Bioanalyzer/Tapestation. Pooled libraries were sequenced on Illumina NextSeq 500 or 1000 platforms (paired-end). Reads were aligned with bowtie2 (settings: –very-sensitive, –no-discordant, –mp 5,1, –np 0, –X 1000), and protein-linked 5′ termini were called using terminalMapper (Perl v5.22.1; ^58^; code at github.com/Neale-Lab). Data were mapped to the Cer3H4L2 *S. cerevisiae* reference and resulting data analysed in R. Primary datasets excluded rDNA, mitochondrial, and 2-µm sequences—except for specific analysis of mitochondrial signals (**Fig. 1d and S2a-j**). Aligned reads per library (**Table S2**) and averaged datasets used for figures (**Table S3**) are reported.

### CC-seq-v2 data analysis: Top3-filtering

Top3 enrichment was achieved by nucleotide-level masking of sites attributable to meiotic Spo11 or Top2. The high-purity Spo11 mask was derived from five *ndt80*Δ *sae2*Δ CC-seq-v2 libraries prepared without ethanol fixation prior to spheroplasting (one each at 2 h and 3 h; three at 4 h), which strongly enriches Spo11 signals. The Top2 mask was built from six ethanol-fixed libraries: three *ndt80*Δ *spo11-yf* and three *ndt80*Δ *spo11-yf top3-md* (2, 3, and 4 h for each genotype). Libraries within each set showed high inter-library correlation. Within a set, candidate Spo11 or Top2 sites were those nucleotide-resolution sites detected in ≥2 replicates on either strand, after accounting for the characteristic top–bottom strand offsets (±1 bp for Spo11; ±3 bp for Top2). Additionally, for each site we computed a score fraction = (mean HpM across replicates) × (replicate detection fraction) and applied set-specific thresholds determined from the empirical distributions: 0.026 for Spo11 and 0.015 for Top2. Replicate-averaged maps were then restricted to sites passing these criteria to yield the final high-purity Spo11 and Top2 masks. To correct strand imbalance from Spo11 concerted cleavage^60^, all 1-bp cognate pairs in the Spo11 mask were equalized to the higher of the two strand signals. For Top3 filtering, CC-seq-v2 datasets were masked sequentially—first by Top2, then by Spo11. Library sizes were rescaled to the total remaining reads after Top2 masking.

### CC-seq-v2 data analysis: Sequence deconvolution

Using high-purity maps of Spo11 and Top2 (see above), together with hotspot-flanking regions derived from the Top3-filtered dataset, we established sequence bias reference profiles for the three proteins (**Fig. S2a**). These reference profiles were then combined in defined proportions (x:y:z) and compared against each unfiltered library. Mixture ratios were systematically varied at 0.1% increments, and the best-fitting combination was identified by root mean square deviation (RMSD) from the observed unfiltered signal.

### CC-seq-v2 data analysis: GC skew

Where indicated in the figure legend, the dotted trace reports a local Top3-enrichment score computed as log₂(–5GC/–5AT). For each position, CC-seq-v2 signal was partitioned by the base 5 nt upstream (−5, in the 5′ orientation of each strand) into two groups: –5GC (G or C) and –5AT (A or T). Each group’s coverage profile was smoothed with a 500-bp Hann window, and the dotted line plots the log₂ ratio of the smoothed –5GC to smoothed –5AT signals.

### CC-seq-v2 data analysis: Averaging signals around loci of interest

Top3-filtered CC-seq-v2 data were aggregated within regions of specified size, centred on the loci of interest, using *CCTools::CCPileup* (**Fig 2c,e; 3c-e; 4c, 5c-d,g**) or *CCTools::CCSum* (**Fig 2d, 5b**), as described previously^1^. The resulting sum total HpM was divided by numbers of loci to give a mean HpM per locus. Convergent and divergent loci were defined as the midpoint between neighbouring genes. Rec8 peaks were defined as described below. Data averaging was either performed at single loci (e.g **Fig. 2e-d; 3d-e; 5b,d,g**), or at scaled intervals between two loci (**Fig. 2c, 3c, 4c, 5c**). Where performed, stratification was based on Rec8 occupancy (**Fig. 3d-e**); sum converging gene expression (**Fig. 2d, 5b**); relative expression of left/right genes (**Fig. 2e**); or—in the case of scaled Divergent–Convergent intervals—the expression level of the intermediary gene, after the exclusion of the bottom 15% of expressed genes (**Fig. 5c**). Gene expression was defined using a previously published RNA-seq dataset employing *P_CUP1_-IME1* induced meiotic SK1 yeast^61^, at the 3 hour timepoint following addition of copper sulphate.

### Analysis of maps of recombination outcomes

Crossover (CO) and noncrossover (NCO) positions were mapped previously^35,37,38^ and reanalysed here. First, CO or NCO event midpoints (**Fig. 5e**) or boundaries (**Fig. 5d,g**) were tabulated in FullMap format (the data format employed by our *CCTools* R package), before averaging around loci of interest, similarly to as for CC-seq-v2 data, using *CCTools::CCPileup* (**Fig. 5d,g**) or *CCTools::CCSum* (**Fig. 5e,f**).

### Calling Rec8 peaks

Rec8 ChIP-seq FASTQ files from wild-type yeast (SRP277061; ^27^) were aligned to the Cer3H4L2 reference using bowtie2. Multimappers were excluded, then reads were de-duplicated using *MACS2::filterdup.* Reads were extended to 200 bp total then replicates were averaged genome-wide. Data were smoothed using a 250-bp sliding Hann window, then signal was computed as IP/input fold enrichment, and finally re-smoothed with a 1-kb window to generate Rec8 occupancy tracks. Peaks were called by identifying local maxima with *CCTools::localMaxima*, then filtering these to include only those with a signal value greater than 1.25.

### Defining Red1 islands

Red1 islands were defined in the Cer3H4L2 genome as described previously^27^. Briefly, Red1 ChIP-seq FASTQ files from *rec8*Δ yeast (SRP277061; ^27^) were aligned to Cer3H4L2 using bowtie2. Multimappers were excluded, then reads were de-duplicated using *MACS2::filterdup.* Reads were extended to 200 bp total then genome-wide signals were calculated as the fold of immunoprecipitated over input coverage within 5 kb bins. Islands were defined as 5 kb bins with a signal score greater or equal to 1.75-fold of the genome-wide average, with adjacent such regions being merged.

### RNA-seq and analysis

RNA-seq was performed as described previously^1^. Briefly, total RNA was extracted from cells using the Monarch Total RNA Miniprep kit, coupled with bead beating using a Fastprep-24 ribolyser. RNA integrity was validated by TBE-agarose gel electrophoresis. Total stranded RNA-seq was performed using the NEBNext Ultra™ II Directional RNA Library Prep Kit for Illumina, coupled with the QIAseq FastSelect – rRNA Yeast Kit. cDNA Libraries were quantified by Bioanalyzer (Agilent), prior to sequencing on the Nextseq 1000 (Illumina). Data were aligned to the Cer3 (R64-1-1) genome using STAR^62^. Gene counts reported by STAR were normalized for library size and composition by the median of ratios method using *DEseq2.* The numbers of aligned reads for each library generated and sequenced in this study are detailed in **Table S2**. Final datasets used for generating figures are detailed in **Table S3**.

## Data availability statement

Datasets analysed in this manuscript are detailed in Supplementary **Table S3**. The CC-seq-v2 and RNA-seq data generated in this study have been deposited in Figshare (https://figshare.com/s/6453d70587876077056f).

## Code availability statement

Custom code used to analyse data reported in this manuscript are available at the following public GitHub repositories at https://github.com/Neale-Lab/CCTools and https://github.com/Neale-Lab/terminalMapper.

## Acknowledgements

T.P. was supported by a Ph.D studentship from the University of Sussex School of Life Sciences. W.H.G. was independently supported by the Biotechnology and Biological Sciences Research Council Discovery Fellowship BB/V005081/1. M.J.N., R.M.A., G.G.B.B. and W.H.G. were supported by the Wellcome Trust Investigator Award 200843/Z/16/Z and Wellcome Trust Discovery Award 225852/Z/22/Z. We thank Antony Oliver (University of Sussex) for the gift of recombinant TDP2 catalytic domain, used in our CC-seq library preparation.

## Author contributions

Conceptualisation: W.H.G.

Data Curation: T.P., W.H.G.

Formal Analysis: T.P., W.H.G.

Funding acquisition: M.J.N., W.H.G.

Investigation: T.P., R.M.A., G.G.B.B., M.J.N., W.H.G.

Methodology: G.G.B.B., M.J.N., W.H.G.

Project administration: M.J.N., W.H.G.

Software: G.G.B.B., W.H.G.

Supervision: M.J.N., W.H.G.

Visualisation: T.P., M.J.N., W.H.G.

Writing—original draft: M.J.N., W.H.G.

Writing—review and editing: M.J.N., W.H.G.

## Supplementary Figure Legends

**Figure S1.**
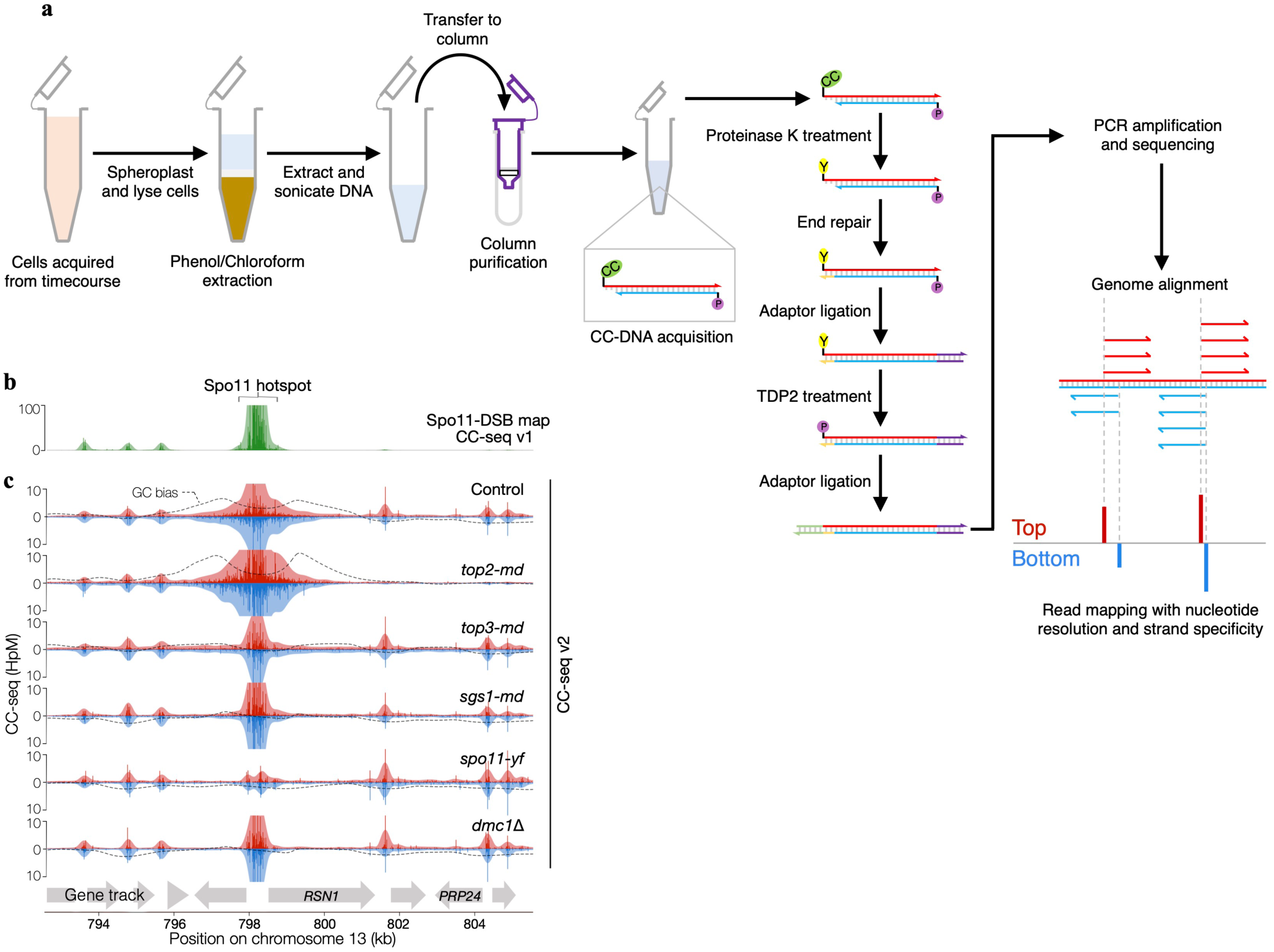
CC-seq v2 maps in meiotic prophase. **a,** Schematic of CC-seq v2, detailing enrichment of protein-DNA covalent complex (CC) molecules, library preparation using recombinant TDP2, and sequencing to reveal positions of CCs with nucleotide resolution and strand specificity. **b**, CC-seq maps over a 20 kb region around a Spo11-DSB hotspot. Green trace, Spo11 DSBs averaged over 2–4 hr in *ndt80*Δ *sae2*Δ cells. Red/blue tracks are unfiltered CC-seq v2 maps of proteins covalently linked to DNA via 5′ phosphotyrosine bonds (Spo11, Top2, Top3) in control (*ndt80*Δ) and mutant strains. Red and blue traces denote top- and bottom-strand 5′ termini.

**Figure S2.**
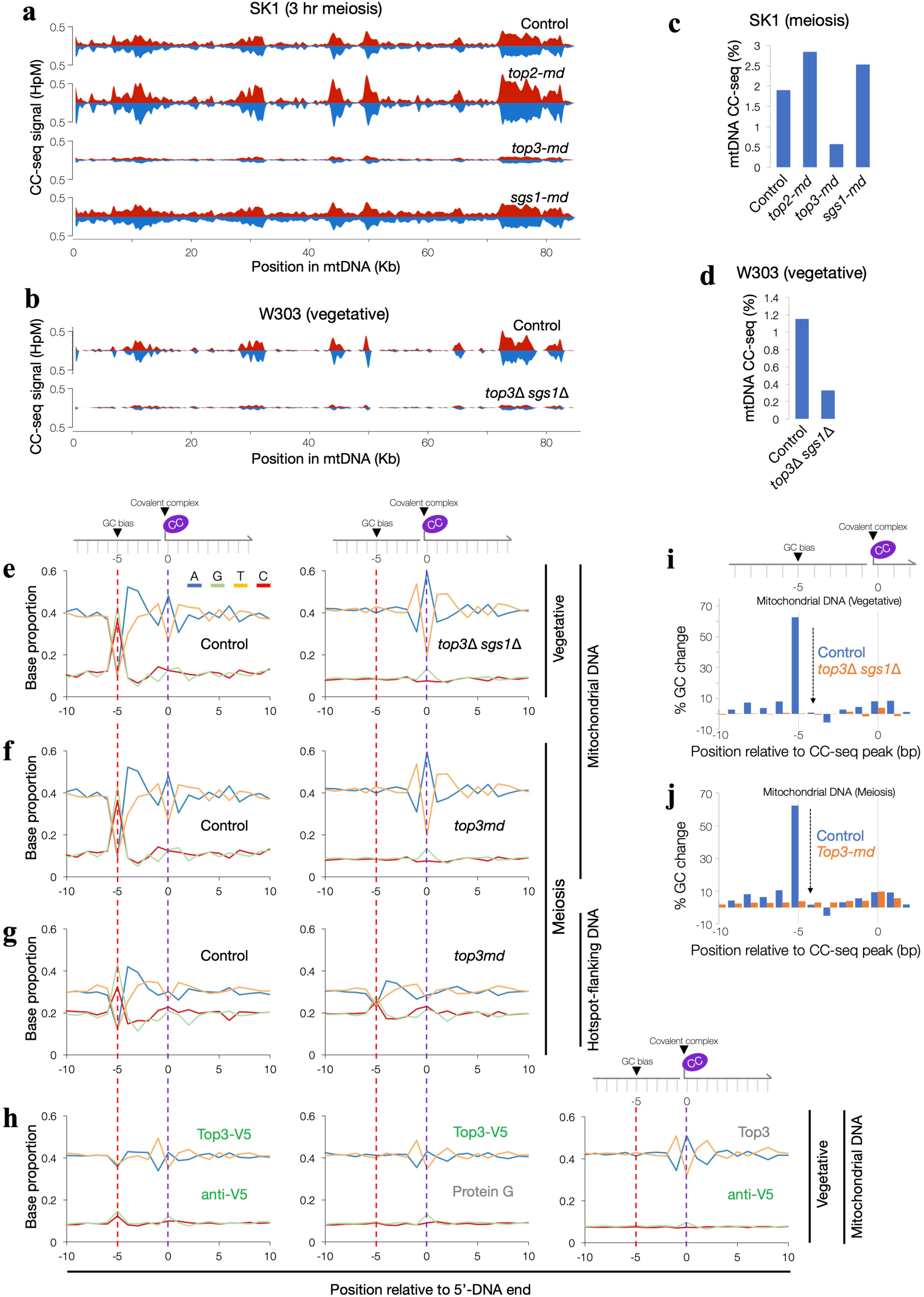
CC-seq maps Top3 activity. **a-b,** Unfiltered CC-seq v2 maps of proteins covalently linked to DNA via 5′ phosphotyrosine bonds within the mitochondria in SK1 *ndt80*Δ strains 6 hr after meiotic induction (**a)** and in asynchronous vegetative W303 control strains at OD_600_ of 2.0 (**b)**. Red and blue traces denote top- and bottom-strand 5′ termini, smoothed with a 1 Kb sliding Hann window. HpM, hits per million mapped reads. **c-d,** The percentage of mitochondrial reads in the CC-seq v2 library, for each of the strains in **a-b**. **e-h,** Base composition over a 20 bp window around unfiltered CC-seq-v2 signal peaks in vegetative W303 mitochondria (**e**), meiotic SK1 mitochondria (**f**), or 500 bp windows flanking Spo11-DSB hotspots (**g**). In **e-g**, left panels are control strains and right panels are *top3*Δ *sgs1*Δ **(e)** or Top3 meiotic-depletion (*top3-md*) cells **(f-g)**. **h,** Base composition around mitochondrial CC-seq signal peaks enriched from a *Top3-V5* strain by anti-V5 immunoprecipitation (left), from the same strain subjected to a negative control immunoprecipitation using protein G (middle), or from an untagged Top3 control strain subjected to immunoprecipitation with anti-V5 immunoprecipitation (right). In **(e-h),** values reported are for the top strand only. Dashed red line highlights position of Top3-dependent GC bias 5 bp upstream of cleavage. Dashed purple line indicates position of covalent complex (log_2_ axis range of ±4). **i–j,** Base composition around unfiltered CC-seq-v2 signal peaks in vegetative W303 mitochondria **(i)**, or meiotic SK1 mitochondria (**j**). Data show percentage change in GC content relative to the local average.

**Figure S3.**
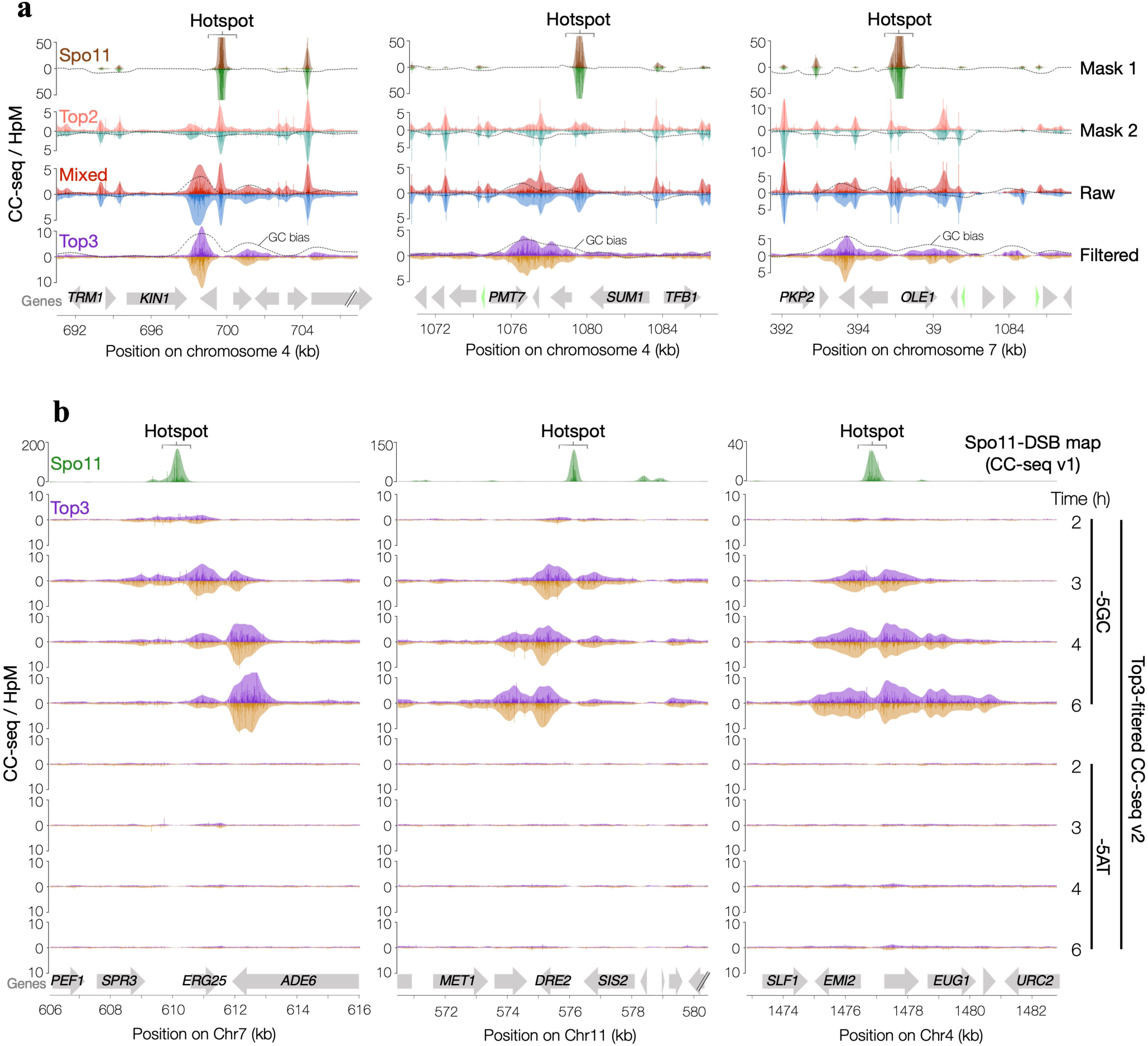
Filtering of Top3 CC-seq signals. **a,** Composite CC-seq maps over a 20 kb region around a Spo11-DSB hotspot. Brown/green tracks are the Spo11 mask generated as described in methods using libraries from 2–4 hr in *ndt80*Δ *sae2*Δ cells. Pink/teal tracks are the Top2 mask generated as described in methods using libraries from 2–4 hr in *ndt80*Δ *spo11-yf* and *ndt80*Δ *spo11-yf top3-md* cells. Red/blue tracks are unfiltered CC-seq v2 maps of proteins covalently linked to DNA via 5′ phosphotyrosine bonds (Spo11, Top2, Top3) in control (*ndt80*Δ) cells. Purple/gold tracks are after Top3-filtering using the Spo11 and Top2 masks. **b**, Top3-filtered CC-seq v2 maps (masked to remove Spo11 and Top2 signals) in control strains. Data were subsequently split into those sites with a - 5GC, or with a -5AT. Top and bottom traces denote top- and bottom-strand 5′ termini. Arrows indicate gene positions.

**Figure S4.**
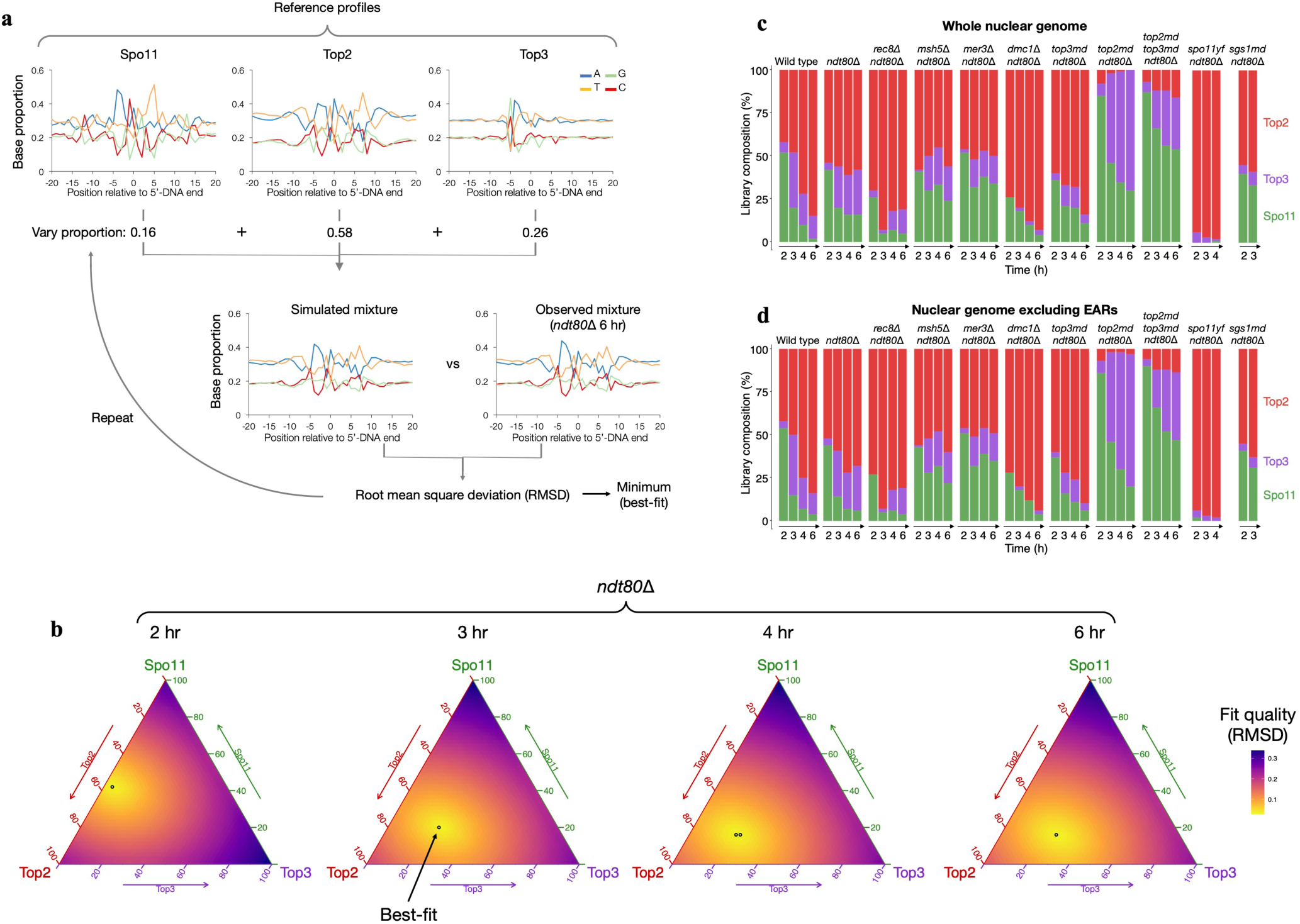
CC-seq signal sequence bias deconvolution. **a,** Schematic of the sequence bias deconvolution algorithm. Reference sequence-bias profiles for Spo11, Top2 and Top3 are presented as base composition over a 20 bp window around CC-seq-v2 peaks. Reference profiles are mixed with a defined proportion to generate a simulated mixture, then compared with the observed mixture in an unfiltered CC-seq-v2 library. Proportions are sampled at 2% increments, and the mixture with the minimal root mean square deviation (RMSD) estimates the component contributions. **b,** Ternary plots of the simulated Spo11:Top2:Top3 proportion vs the RMSD relative to the observed unfiltered CC-seq-v2 library for timepoints through meiosis in *ndt80*Δ cells. Best fits are indicated by black circles. **c-d,** Stacked bar showing best-fit estimates for the relative fraction of Top2 (red), Spo11 (green) and Top3 (purple) signal at each timepoint genome-wide **(c)** or after excluded the chromosome end-adjacent regions; the small chromosomes 1, 3 and 6; and the rDNA-containing chromosome 12 (**d**).

**Figure S5.**
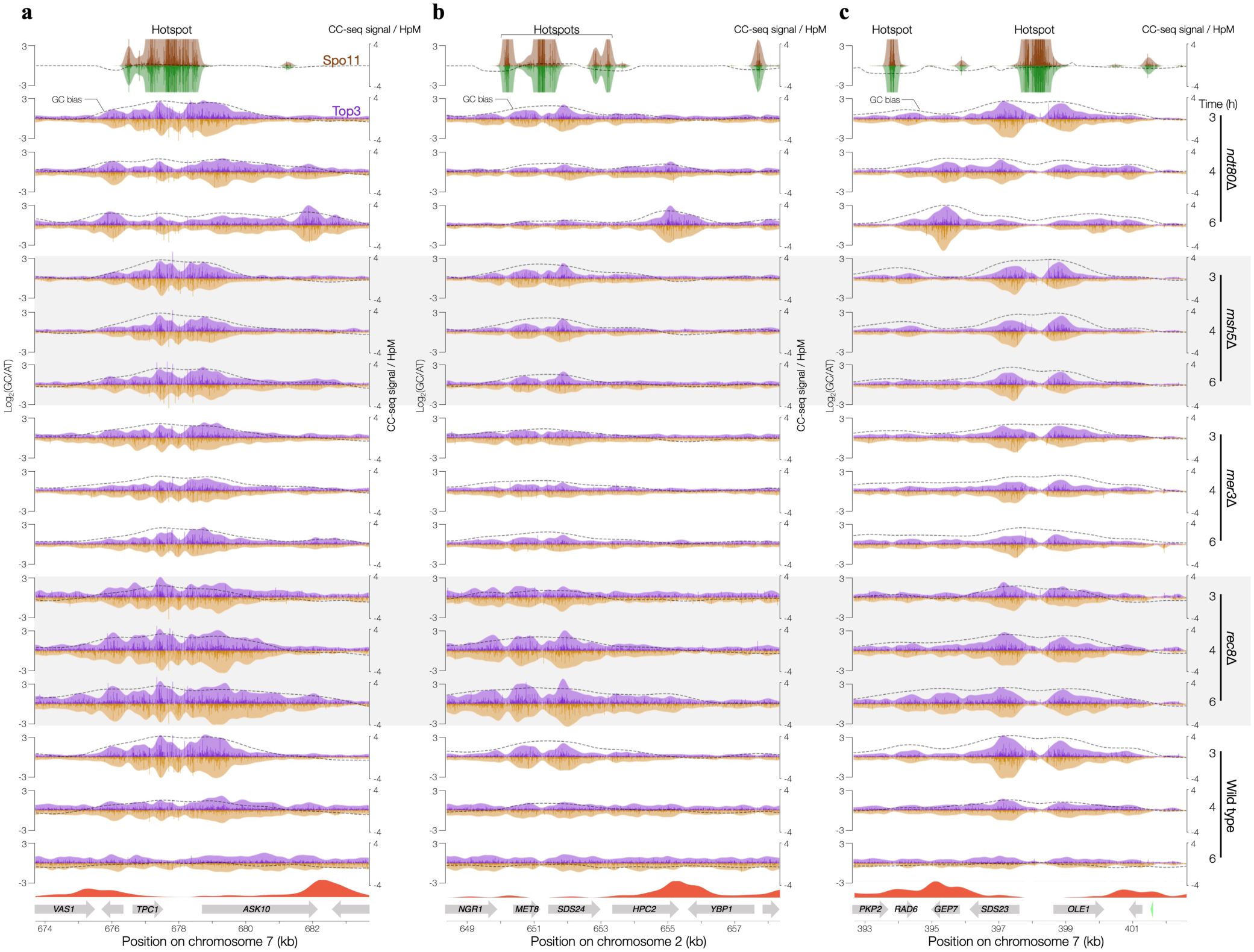
Top3 migration in wild type, *ndt80*Δ, *ndt80*Δ *mer3*Δ, *ndt80*Δ *msh5*Δ and *ndt80*Δ *rec8*Δ. **a-b,** Composite maps of strand-specific Spo11 activity (CC-seq v1, 2–4 h, *ndt80Δ sae2Δ* and *ndt80Δ sae2Δ rec8*Δ, brown/green), Top3 activity (Top3-filtered CC-seq v2, 3 h, 4 h and 6 h, purple/gold), and Rec8 occupancy (ChIP-seq, 3 h, orange), at three 10 kb regions. Lighter tones show 500 bp Hann smoothing; dashed lines mark skews towards GC bases 5 bp upstream of cleavage. Grey arrows mark genes. HpM, hits per million mapped reads.

**Figure S6.**
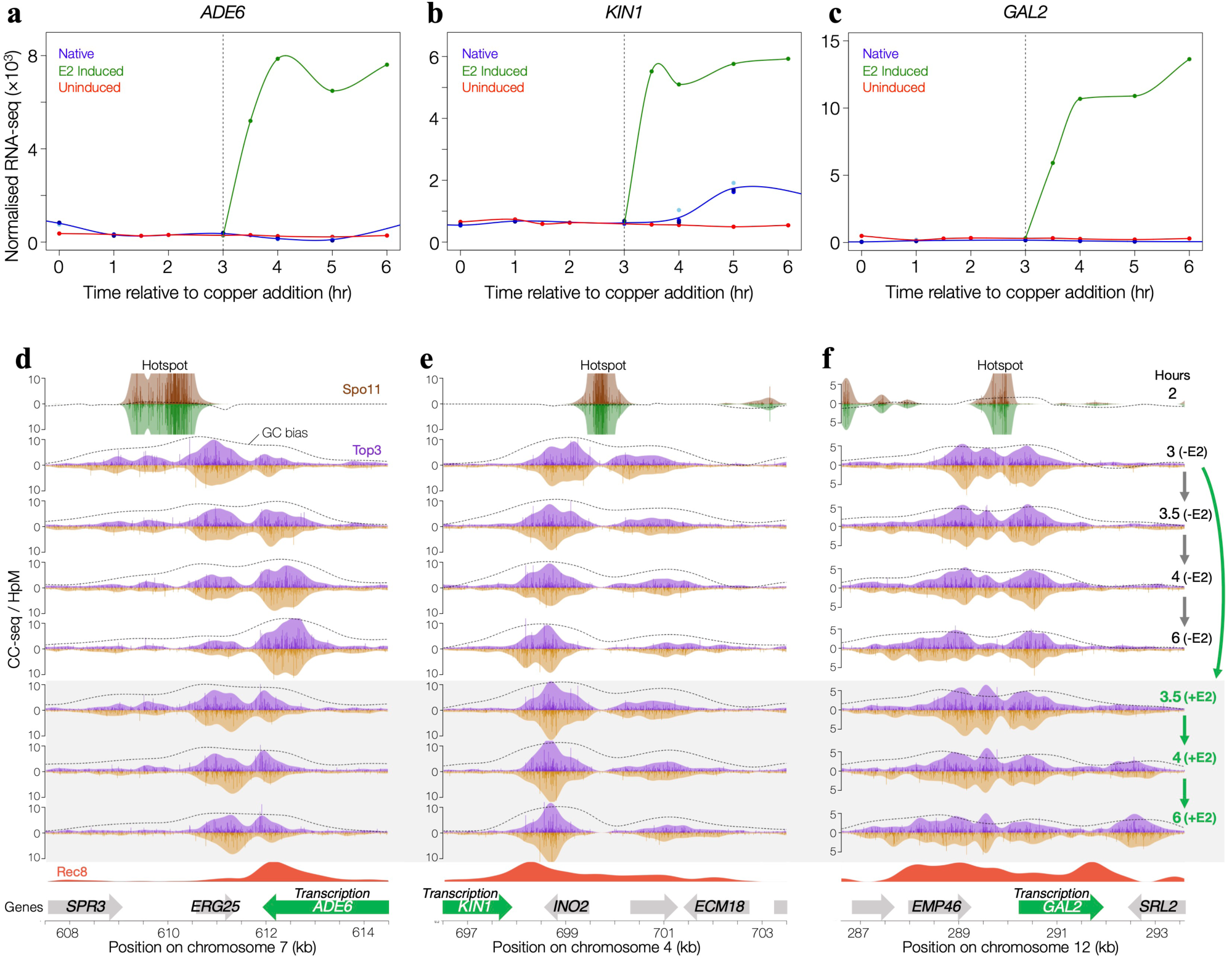
Top3 migration is patterned by active transcription. **a-c,** RNA-seq read counts for genes: *ADE6* **(a)***, KIN1* **(b)**, and *GAL2* **(c),** at timepoints following meiotic induction with copper. Red and green lines are for *ndt80Δ GAL4::ER P_GAL1_ADE6 pGAL-KIN1* cells uninduced (red) or induced (blue) with estradiol (E2) at the 3 hr time point. Blue lines are for a previously published data set with unmodified *ADE6* and *KIN1* loci and no estradiol^61^. All strains contain the *P_CUP1_IME1* gene. Read counts are normalized by the median of ratios method using the *DEseq2* R package^63^. **d-f**, Composite maps at estradiol-inducible loci (*P_GAL1_ADE6, P_GAL1_KIN1* and *GAL2*) in *ndt80Δ GAL4::ER P_GAL1_ADE6 P_GAL1_KIN1* cells. Top3 activity (purple/gold, 3 hr meiosis induction ±30 min, ±1 hr ±3 hr estradiol induction), and reference tracks of Spo11 activity (brown/green) and Rec8 occupancy (orange; ^17^) from control cells are shown with 500 bp Hann smoothing. Dashed lines mark 5’ GC skews; grey arrows mark genes; green arrows indicate induced transcription. HpM, hits per million mapped reads.

**Figure S7.**
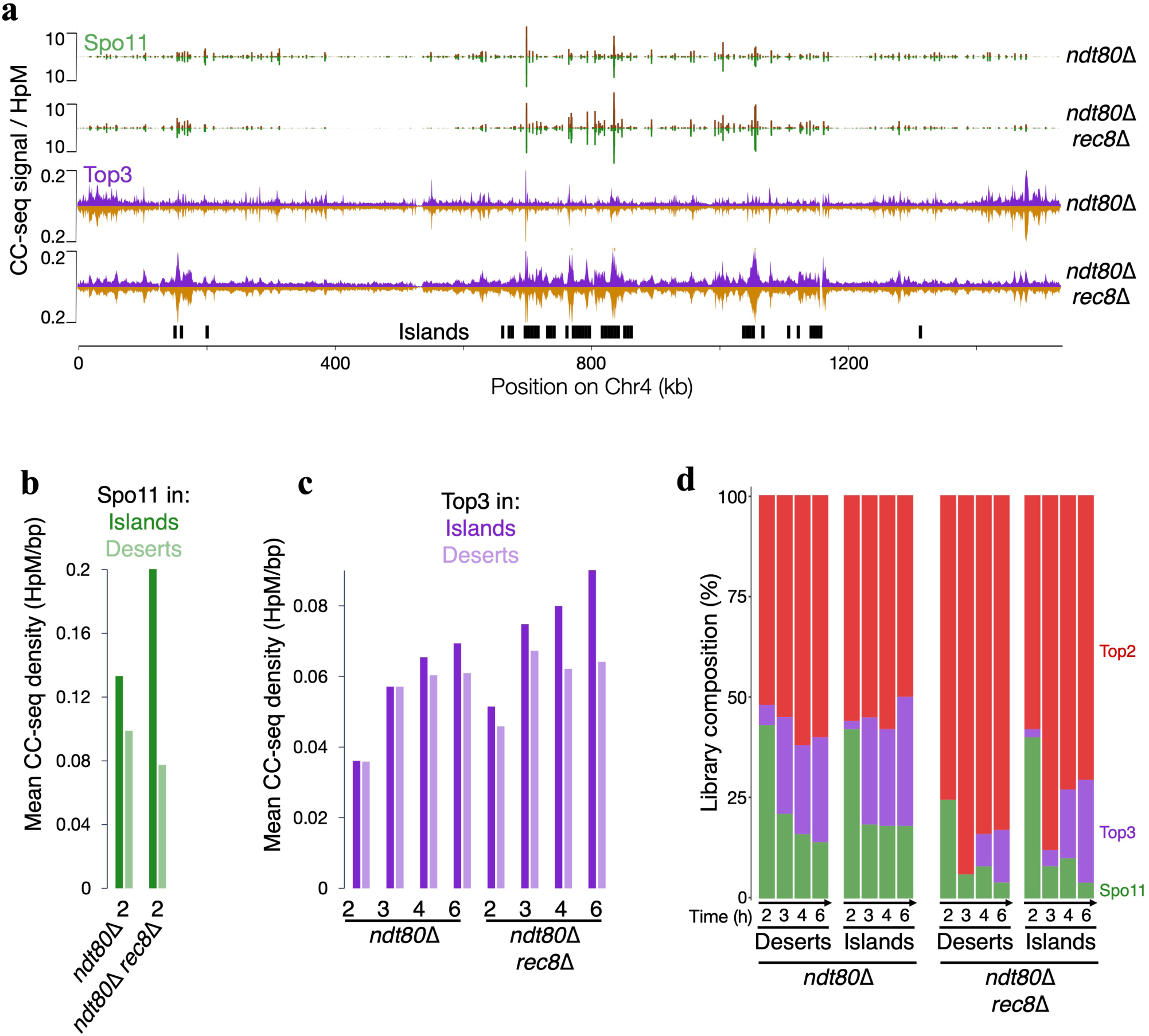
Cohesin influences Spo11 and Top3 activity genome-wide. **a,** Composite map of strand-specific Spo11 activity in *ndt80Δ sae2Δ* and *ndt80Δ sae2Δ rec8*Δ cells (CC-seq, brown/green), Top3 activity in *ndt80Δ* and *ndt80Δ rec8*Δ cells, (Top3-filtered, CC-seq v2, 6 h, purple/gold), and Red1 “island regions” (3 h, black rectangles), across chromosome 4. HpM, hits per million mapped reads. **b-c,** barplots of mean density of Spo11 CC-seq (**b**), or Top3-filtered CC-seq (**c**) signals within islands (dark green) and desert regions (light green), in *ndt80Δ* and *ndt80Δ rec8Δ* cells. **d**, Stacked bar showing best-fit estimates for the relative fraction of Top2 (red), Spo11 (green) and Top3 (purple) signal at each timepoint (see Methods), within islands and desert regions.

**Supplementary Table 1.**
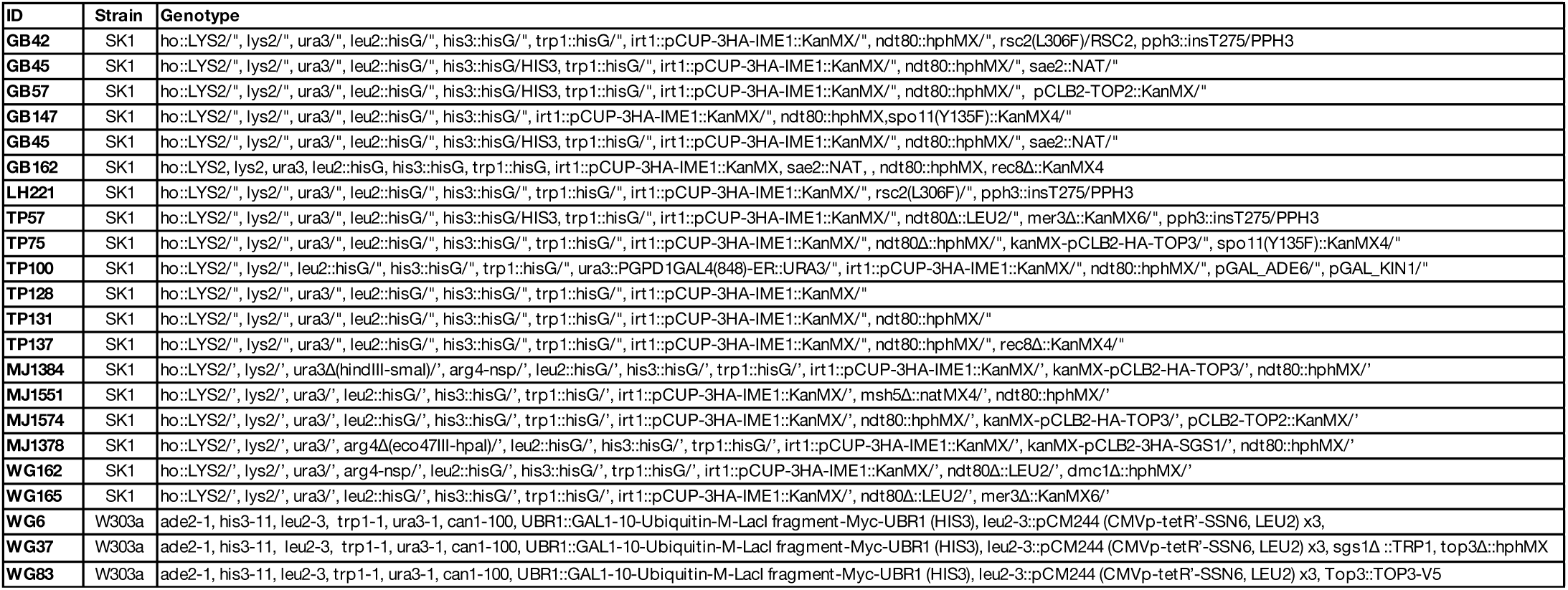
S. cerevisiae strains used in this study. All strains are from the SK1 or W303 background.

**Supplementary Table 2.**
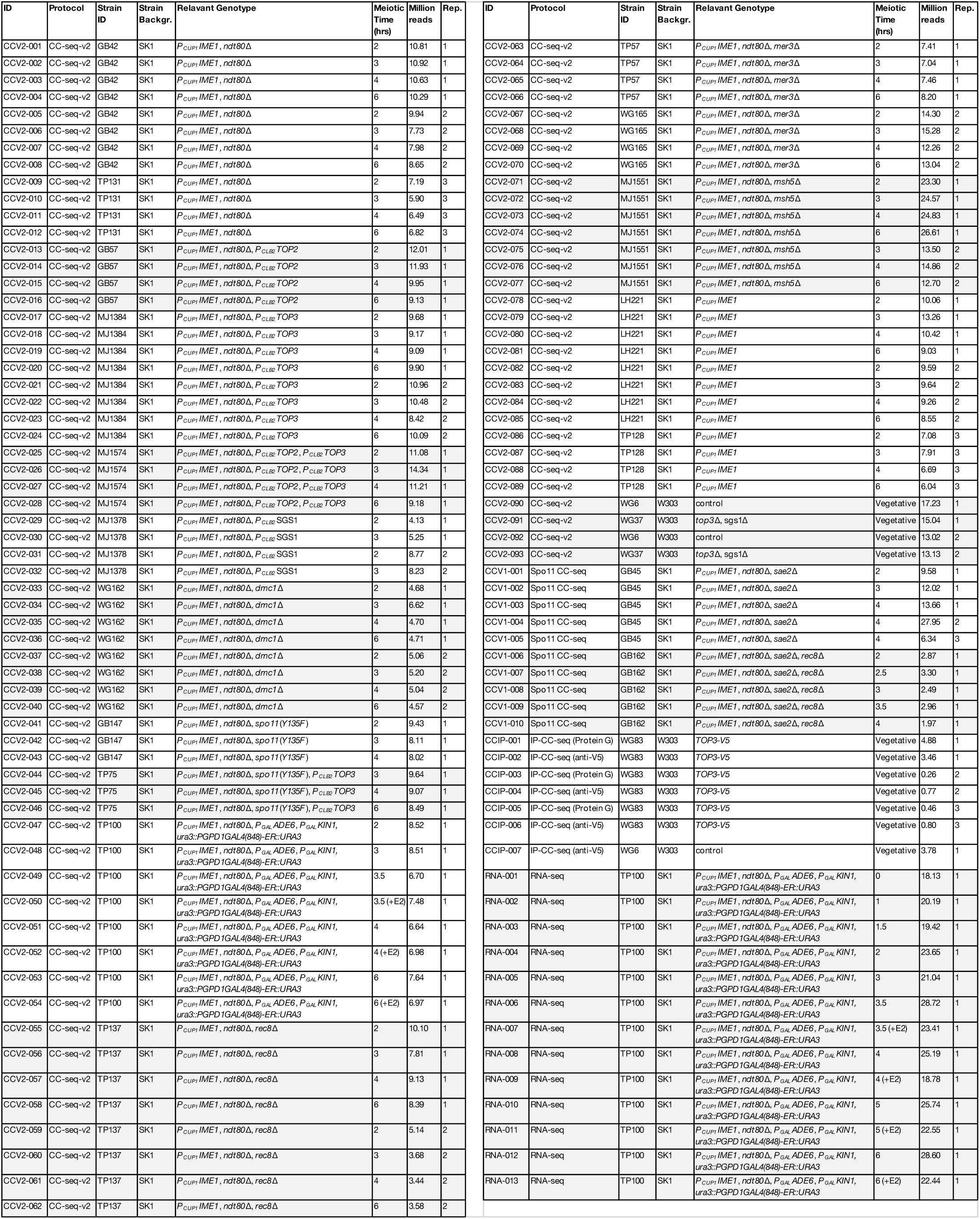
CC-seq-v2, CC-seq-v2, IP-CC-seq, and RNA-seq datasets generated in this study. Individual libraries were prepared as indicated and sequenced using paired-end Illumina sequencing to the indicated mapped read depth.

**Supplementary Table 3.**
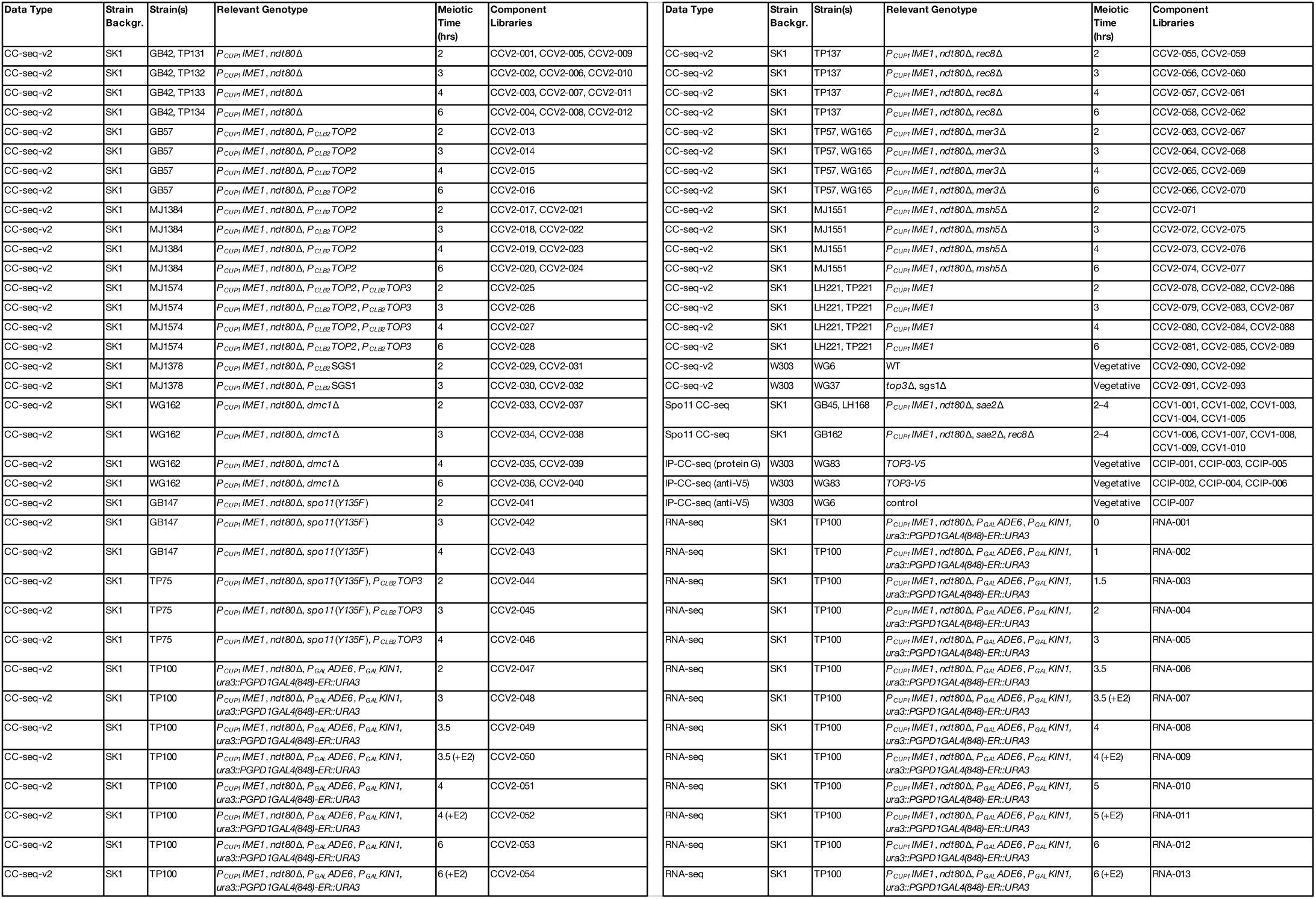
Final dataset averages used in this study. Component CC-seq-v2 libraries were averaged together with equal weighting to generate the final datasets indicated.

## REFERENCES

1 Gittens, W. H., Allison, R. M., Wright, E. M., Brown, G. G. B. & Neale, M. J. Osmotic disruption of chromatin induces Topoisomerase 2 activity at sites of transcriptional stress. Nat Commun 15, 10606, doi:10.1038/s41467-024-54567-6 (2024).

2 Keeney, S., Giroux, C. N. & Kleckner, N. Meiosis-specific DNA double-strand breaks are catalyzed by Spo11, a member of a widely conserved protein family. Cell 88, 375–384 (1997).

3 Wang, J. C. Cellular roles of DNA topoisomerases: a molecular perspective. Nat Rev Mol Cell Biol 3, 430–440 (2002).

4 Gangloff, S., de Massy, B., Arthur, L., Rothstein, R. & Fabre, F. The essential role of yeast topoisomerase III in meiosis depends on recombination. The EMBO Journal 18, 1701–1711, 10.1093/emboj/18.6.1701 (1999).

5 Norman-Axelsson, U., Durand-Dubief, M., Prasad, P. & Ekwall, K. DNA Topoisomerase III Localizes to Centromeres and Affects Centromeric CENP-A Levels in Fission Yeast. PLOS Genetics 9, e1003371, doi:10.1371/journal.pgen.1003371 (2013).

6 Kaur, H., De Muyt, A. & Lichten, M. Top3-Rmi1 DNA Single-Strand Decatenase Is Integral to the Formation and Resolution of Meiotic Recombination Intermediates. Molecular Cell 57, 583–594, 10.1016/j.molcel.2015.01.020 (2015).

7 Tang, S., Wu, M. K. Y., Zhang, R. & Hunter, N. Pervasive and essential roles of the Top3-Rmi1 decatenase orchestrate recombination and facilitate chromosome segregation in meiosis. Mol Cell 57, 607–621, doi:10.1016/j.molcel.2015.01.021 (2015).

8 Bergerat, A. et al. An atypical topoisomerase II from archaea with implications for meiotic recombination. Nature 386, 414–417, doi:10.1038/386414a0 (1997).

9 Bishop, D. K., Park, D., Xu, L. & Kleckner, N. DMC1: A meiosis-specific yeast homolog of E. coli recA required for recombination, synaptonemal complex formation, and cell cycle progression. Cell 69, 439–456, 10.1016/0092-8674(92)90446-J (1992).

10 Wang, Y., Lyu, Y. L. & Wang, J. C. Dual localization of human DNA topoisomerase IIIα to mitochondria and nucleus. Proceedings of the National Academy of Sciences 99, 12114–12119, doi:doi:10.1073/pnas.192449499 (2002).

11 Nicholls, T. J. et al. Topoisomerase 3α Is Required for Decatenation and Segregation of Human mtDNA. Molecular Cell 69, 9–23.e26, 10.1016/j.molcel.2017.11.033 (2018).

12 Menger, K. E. et al. Two type I topoisomerases maintain DNA topology in human mitochondria. Nucleic Acids Res 50, 11154–11174, doi:10.1093/nar/gkac857 (2022).

13 Wu, J., Feng, L. & Hsieh, T.-s. Drosophila topo IIIα is required for the maintenance of mitochondrial genome and male germ-line stem cells. Proceedings of the National Academy of Sciences 107, 6228–6233, doi:10.1073/pnas.1001855107 (2010).

14 Kim, R. A. & Wang, J. C. Identification of the yeast TOP3 gene product as a single strand-specific DNA topoisomerase. Journal of Biological Chemistry 267, 17178–17185, 10.1016/S0021-9258(18)41910-2 (1992).

15 Xu, L., Ajimura, M., Padmore, R., Klein, C. & Kleckner, N. NDT80, a meiosis-specific gene required for exit from pachytene in Saccharomyces cerevisiae. Mol Cell Biol 15, 6572–6581, doi:10.1128/mcb.15.12.6572 (1995).

16 Allers, T. & Lichten, M. Differential timing and control of noncrossover and crossover recombination during meiosis. Cell 106, 47–57, doi:10.1016/s0092-8674(01)00416-0 (2001).

17 Panizza, S. et al. Spo11-accessory proteins link double-strand break sites to the chromosome axis in early meiotic recombination. Cell 146, 372–383, doi:10.1016/j.cell.2011.07.003 (2011).

18 Klein, F. et al. A Central Role for Cohesins in Sister Chromatid Cohesion, Formation of Axial Elements, and Recombination during Yeast Meiosis. Cell 98, 91–103, doi:10.1016/S0092-8674(00)80609-1 (1999).

19 Schalbetter, S. A., Fudenberg, G., Baxter, J., Pollard, K. S. & Neale, M. J. Principles of meiotic chromosome assembly revealed in S. cerevisiae. Nat Commun 10, 4795, doi:10.1038/s41467-019-12629-0 (2019).

20 Sun, X. et al. Transcription dynamically patterns the meiotic chromosome-axis interface. eLife 4, e07424, doi:10.7554/eLife.07424 (2015).

21 Blat, Y. & Kleckner, N. Cohesins bind to preferential sites along yeast chromosome III, with differential regulation along arms versus the centric region. Cell 98, 249–259, doi:10.1016/s0092-8674(00)81019-3 (1999).

22 Kim, K. P. et al. Sister Cohesion and Structural Axis Components Mediate Homolog Bias of Meiotic Recombination. Cell 143, 924–937, 10.1016/j.cell.2010.11.015 (2010).

23 Davidson, I. F. et al. DNA loop extrusion by human cohesin. Science 366, 1338–1345, doi:10.1126/science.aaz3418 (2019).

24 Kim, Y., Shi, Z., Zhang, H., Finkelstein, I. J. & Yu, H. Human cohesin compacts DNA by loop extrusion. Science 366, 1345–1349, doi:10.1126/science.aaz4475 (2019).

25 Fudenberg, G. et al. Formation of Chromosomal Domains by Loop Extrusion. Cell Reports 15, 2038–2049, 10.1016/j.celrep.2016.04.085 (2016).

26 Lengronne, A. et al. Cohesin relocation from sites of chromosomal loading to places of convergent transcription. Nature 430, 573–578, doi:10.1038/nature02742 (2004).

27 Heldrich, J. et al. Two pathways drive meiotic chromosome axis assembly in Saccharomyces cerevisiae. Nucleic Acids Res 50, 4545–4556, doi:10.1093/nar/gkac227 (2022).

28 Börner, G. V., Kleckner, N. & Hunter, N. Crossover/Noncrossover Differentiation, Synaptonemal Complex Formation, and Regulatory Surveillance at the Leptotene/Zygotene Transition of Meiosis. Cell 117, 29–45, 10.1016/S0092-8674(04)00292-2 (2004).

29 Snowden, T., Acharya, S., Butz, C., Berardini, M. & Fishel, R. hMSH4-hMSH5 Recognizes Holliday Junctions and Forms a Meiosis-Specific Sliding Clamp that Embraces Homologous Chromosomes. Molecular Cell 15, 437–451, 10.1016/j.molcel.2004.06.040 (2004).

30 Nakagawa, T. & Kolodner, R. D. Saccharomyces cerevisiae Mer3 Is a DNA Helicase Involved in Meiotic Crossing Over. Molecular and Cellular Biology 22, 3281–3291, doi:10.1128/MCB.22.10.3281-3291.2002 (2002).

31 Mazina, O. M., Mazin, A. V., Nakagawa, T., Kolodner, R. D. & Kowalczykowski, S. C. Saccharomyces cerevisiae Mer3 Helicase Stimulates 3′–5′ Heteroduplex Extension by Rad51: Implications for Crossover Control in Meiotic Recombination. Cell 117, 47–56, 10.1016/S0092-8674(04)00294-6 (2004).

32 Subramanian, V. V. et al. Persistent DNA-break potential near telomeres increases initiation of meiotic recombination on short chromosomes. Nature Communications 10, 970, doi:10.1038/s41467-019-08875-x (2019).

33 Schwacha, A. & Kleckner, N. Identification of double Holliday junctions as intermediates in meiotic recombination. Cell 83, 783–791, doi:10.1016/0092-8674(95)90191-4 (1995).

34 Hunter, N. & Kleckner, N. The Single-End Invasion: An Asymmetric Intermediate at the Double-Strand Break to Double-Holliday Junction Transition of Meiotic Recombination. Cell 106, 59–70, doi:10.1016/S0092-8674(01)00430-5 (2001).

35 Crawford, M. R. et al. Separable roles of the DNA damage response kinase Mec1ATR and its activator Rad24RAD17 during meiotic recombination. PLOS Genetics 20, e1011485, doi:10.1371/journal.pgen.1011485 (2024).

36 Benjamin, K. R., Zhang, C., Shokat, K. M. & Herskowitz, I. Control of landmark events in meiosis by the CDK Cdc28 and the meiosis-specific kinase Ime2. Genes & Development 17, 1524–1539 (2003).

37 Mancera, E., Bourgon, R., Brozzi, A., Huber, W. & Steinmetz, L. M. High-resolution mapping of meiotic crossovers and non-crossovers in yeast. Nature 454, 479–485, doi:10.1038/nature07135 (2008).

38 Oke, A., Anderson, C. M., Yam, P. & Fung, J. C. Controlling Meiotic Recombinational Repair – Specifying the Roles of ZMMs, Sgs1 and Mus81/Mms4 in Crossover Formation. PLOS Genetics 10, e1004690, doi:10.1371/journal.pgen.1004690 (2014).

39 Henggeler, A., Orlić, L., Velikov, D. & Matos, J. Holliday junction–ZMM protein feedback enables meiotic crossover assurance. Nature 647, 766–775, doi:10.1038/s41586-025-09559-x (2025).

40 Tang, S. et al. Protecting double Holliday junctions ensures crossing over during meiosis. Nature 647, 776–785, doi:10.1038/s41586-025-09555-1 (2025).

41 Ahuja, J. S., Harvey, C. S., Wheeler, D. L. & Lichten, M. Repeated strand invasion and extensive branch migration are hallmarks of meiotic recombination. Molecular Cell 81, 4258–4270.e4254, 10.1016/j.molcel.2021.08.003 (2021).

42 Chen, S. H., Plank, J. L., Willcox, S., Griffith, J. D. & Hsieh, T. S. Top3α is required during the convergent migration step of double Holliday junction dissolution. PLoS One 9, e83582, doi:10.1371/journal.pone.0083582 (2014).

43 Cejka, P., Plank, J. L., Bachrati, C. Z., Hickson, I. D. & Kowalczykowski, S. C. Rmi1 stimulates decatenation of double Holliday junctions during dissolution by Sgs1–Top3. Nature Structural & Molecular Biology 17, 1377–1382, doi:10.1038/nsmb.1919 (2010).

44 Altmannova, V. et al. Biochemical characterisation of Mer3 helicase interactions and the protection of meiotic recombination intermediates. Nucleic Acids Res 51, 4363–4384, doi:10.1093/nar/gkad175 (2023).

45 Sanchez, A. et al. Exo1 recruits Cdc5 polo kinase to MutLγ to ensure efficient meiotic crossover formation. Proceedings of the National Academy of Sciences of the United States of America 117, 30577–30588, doi:10.1073/pnas.2013012117 (2020).

46 Marsolier-Kergoat, M. C., Khan, M. M., Schott, J., Zhu, X. & Llorente, B. Mechanistic View and Genetic Control of DNA Recombination during Meiosis. Mol Cell 70, 9–20.e26, doi:10.1016/j.molcel.2018.02.032 (2018).

47 Thacker, D., Mohibullah, N., Zhu, X. & Keeney, S. Homologue engagement controls meiotic DNA break number and distribution. Nature 510, 241–246, doi:10.1038/nature13120 (2014).

48 Orlić, L., Henggeler, A., Nagy, J. & Matos, J. Distinct and overlapping roles of MutLγ, Mus81-Mms4, and STR in meiotic Holliday junction processing. Research Square, 10.21203/rs.3.rs-8593203/v1 (2026).

49 Pipathsouk, A., Belotserkovskii, B. P. & Hanawalt, P. C. When transcription goes on Holliday: Double Holliday junctions block RNA polymerase II transcription in vitro. Biochimica et Biophysica Acta (BBA) - Gene Regulatory Mechanisms 1860, 282–288, 10.1016/j.bbagrm.2016.12.002 (2017).

50 Forth, S., Deufel, C., Patel, Smita S. & Wang, Michelle D. Direct Measurements of Torque During Holliday Junction Migration. Biophysical Journal 101, L5–L7, doi:10.1016/j.bpj.2011.05.066 (2011).

51 Zhang, H. L., Malpure, S. & DiGate, R. J. Escherichia coli DNA Topoisomerase III Is a Site-specific DNA Binding Protein That Binds Asymmetrically to Its Cleavage Site (∗). Journal of Biological Chemistry 270, 23700–23705, 10.1074/jbc.270.40.23700 (1995).

52 Goulaouic, H. et al. Purification and characterization of human DNA topoisomerase IIIα. Nucleic Acids Research 27, 2443–2450, doi:10.1093/nar/27.12.2443 (1999).

53 Yang, X., Saha, S., Yang, W., Neuman, K. C. & Pommier, Y. Structural and biochemical basis for DNA and RNA catalysis by human Topoisomerase 3β. Nature Communications 13, 4656, doi:10.1038/s41467-022-32221-3 (2022).

54 Yang, X., Chen, X., Yang, W. & Pommier, Y. Structural insights into human topoisomerase 3β DNA and RNA catalysis and nucleic acid gate dynamics. Nature Communications 16, 834, doi:10.1038/s41467-025-55959-y (2025).

55 Kane, S. M. & Roth, R. Carbohydrate metabolism during ascospore development in yeast. J Bacteriol 118, 8–14, doi:10.1128/jb.118.1.8-14.1974 (1974).

56 Chia, M. & van Werven, F. J. Temporal Expression of a Master Regulator Drives Synchronous Sporulation in Budding Yeast. G3 (Bethesda) 6, 3553-3560, doi:10.1534/g3.116.034983 (2016).

57 Brown, G. G. B., Gittens, W. H., Allison, R. M., Oliver, A. W. & Neale, M. J. CC-seq: Nucleotide-Resolution Mapping of Spo11 DNA Double-Strand Breaks in S. cerevisiae Cells. Methods in molecular biology (Clifton, N.J.) 2818, 3–22, doi:10.1007/978-1-0716-3906-1_1 (2024).

58 Gittens, W. H. et al. A nucleotide resolution map of Top2-linked DNA breaks in the yeast and human genome. Nat Commun 10, 4846, doi:10.1038/s41467-019-12802-5 (2019).

59 Hornyak, P. et al. Mode of action of DNA-competitive small molecule inhibitors of tyrosyl DNA phosphodiesterase 2. Biochem J 473, 1869–1879, doi:10.1042/bcj20160180 (2016).

60 Johnson, D. et al. Concerted cutting by Spo11 illuminates meiotic DNA break mechanics. Nature 594, 572–576, doi:10.1038/s41586-021-03389-3 (2021).

61 Scutenaire, J. et al. The S. cerevisiae m6A-reader Pho92 promotes timely meiotic recombination by controlling key methylated transcripts. Nucleic Acids Res 51, 517–535, doi:10.1093/nar/gkac640 (2023).

62 Dobin, A. et al. STAR: ultrafast universal RNA-seq aligner. Bioinformatics 29, 15–21, doi:10.1093/bioinformatics/bts635 (2013).

63 Love, M. I., Huber, W. & Anders, S. Moderated estimation of fold change and dispersion for RNA-seq data with DESeq2. Genome Biology 15, 550, doi:10.1186/s13059-014-0550-8 (2014).

